# Endogenous tagging using split mNeonGreen in human iPSCs for live imaging studies

**DOI:** 10.1101/2023.06.28.546942

**Authors:** Mathieu C. Husser, Nhat P. Pham, Chris Law, Flavia R. B. Araujo, Vincent J.J. Martin, Alisa Piekny

## Abstract

Endogenous tags have become invaluable tools to visualize and study native proteins in live cells. However, generating human cell lines carrying endogenous tags is difficult due to the low efficiency of homology-directed repair. Recently, an engineered split mNeonGreen protein was used to generate a large-scale endogenous tag library in HEK293 cells. Using split mNeonGreen for large-scale endogenous tagging in human iPSCs would open the door to studying protein function in healthy cells and across differentiated cell types. We engineered an iPS cell line to express the large fragment of the split mNeonGreen protein (mNG2_1-10_) and showed that it enables fast and efficient endogenous tagging of proteins with the short fragment (mNG2_11_). We also demonstrate that neural network-based image restoration enables live imaging studies of highly dynamic cellular processes such as cytokinesis in iPSCs. This work represents the first step towards a genome-wide endogenous tag library in human stem cells.

## Introduction

Since GFP was first described as a fluorescent reporter (Chalfie et al., 1994), the use of fluorescent proteins has become a standard method to study the localization and function of proteins inside living cells. Generally, reporter protein fusions are transiently or stably expressed from exogenous plasmids carrying their own promoter. However, expression levels can be quite high and variable from these strong, non-specific promoters (Husser et al., 2022). Further, over-expression can induce artefacts of localization and protein-protein interactions, making data challenging to interpret (Gibson et al., 2013; Mahen et al., 2014). With the advent of gene editing tools such as CRISPR/Cas9, the genes encoding fluorescent proteins can now be inserted directly into the genome at a desired locus (Bukhari & Muller, 2019; Husser et al., 2021). CRISPR/Cas9 is used to introduce a double-stranded break at the target site, which can be repaired by homology directed repair (HDR) using a repair template carrying the fluorescent marker (Verma et al., 2017). With this approach, the protein of interest is still expressed from its endogenous promoter, fused with the fluorescent protein. This enables the study of proteins at endogenous expression levels and provides more reliable measurements of protein behavior (Dambournet et al., 2018; Doyon et al., 2011; Husser et al., 2022; Mahen et al., 2014). Endogenous tagging is commonly done in model organisms such as *Saccharomyces cerevisiae*, for which whole-genome libraries of endogenous tags have been generated (Huh et al., 2003). However, gene editing in human cells is less widely used due to limitations in efficiency caused by transfection and HDR, among other issues. Because of these bottlenecks, the majority of human proteins have not been tagged and studied at the endogenous level. Although several efforts have been made to tag multiple proteins endogenously in various human cell lines (*e.g.* Husser et al., 2022; Roberts et al., 2017), the generation of a genome-wide library of tagged proteins in human cells requires higher throughput.

The recently developed self-complementing split fluorescent proteins can be used as tools for large-scale endogenous tagging (Feng et al., 2017; Feng et al., 2019; Kamiyama et al., 2016; Tamura et al., 2021; Zhou et al., 2020). In the split mNeonGreen system, the mNeonGreen protein is expressed as two separate fragments: a large fragment composed of the first ten beta-strands of mNeonGreen (mNG2_1-10_) and a short fragment corresponding to the eleventh beta strand (mNG211; Feng et al., 2017). The two fragments have been engineered to form a functional fluorescent protein when co-expressed (Feng et al., 2017). This interaction is characterized by a high complementation efficiency and is irreversible once the reconstituted protein has folded (Feng et al., 2019; Koker et al., 2018). Moreover, the split mNeonGreen2 protein retains most of the brightness of the original mNeonGreen (Feng et al., 2017). This system allows for easy and efficient endogenous tagging with mNG2_11_ in cells where mNG2_1-10_ is constitutively expressed (Cho et al., 2022; Leonetti et al., 2016; Mahdessian et al., 2021). The mNG2_11_ fragment is only 16 amino acids long, so it can be inserted into the genome by HDR using a repair template with short homology arms (generally 40-80 bp), which can be purchased commercially as a single-stranded oligo-deoxynucleotide (ssODN). This enables large-scale tagging using commercially synthesized ssODNs and sgRNAs to target multiple proteins in parallel (Cho et al., 2022; Leonetti et al., 2016). However, this approach requires the generation of a parental cell line that constitutively expresses the mNG2_1-10_ fragment. This has been done by random lentiviral integration followed by antibiotic selection, which is fast but generates a heterogeneous population where the expression of the large fragment is inconsistent across the cell population and is subject to epigenetic silencing (Cabrera et al., 2022). Large-scale endogenous tagging in HEK293 cells was recently achieved by the OpenCell project, with 1,310 proteins tagged to date (Cho et al., 2022). However, while this library will be a valuable resource for the community, there is a need to study protein function in other cell types, where the mechanisms regulating biological processes could vary drastically.

Since their discovery in 2007, human induced pluripotent stem cells (iPSCs) have become a popular tool to study human cells in developmental contexts and to identify disease-causing mutations, amongst other valuable applications (Shi et al., 2017; Takahashi et al., 2007). iPSCs are derived from somatic cells that are reprogrammed to be self-renewing and pluripotent. These cells can be cultured and differentiated into any desired cell type *in vitro* using specific protocols (*e.g.* Grancharova et al., 2021; Hong & Do, 2019; Oceguera-Yanez et al., 2022). Despite this incredible resource, few studies have investigated cellular processes in human stem cells and differentiated cell types with high spatiotemporal resolution (Dambournet et al., 2018; Viana et al., 2023). For example, several studies describe the regulation of mitosis and cytokinesis in mouse embryos and embryonic stem cells (Chaigne et al., 2020; Chaigne et al., 2021; Paim & FitzHarris, 2022), but not in human stem cells. Since most of our knowledge of human cell cytokinesis is derived from diseased and/or transformed cell lines, the field will benefit greatly from studying iPSCs before and after differentiation in an isogenic context, particularly since they represent ‘healthy’ human cells (Dambournet et al., 2018; Drubin & Hyman, 2017).

Here, we used the split mNeonGreen system for endogenous tagging in human iPSCs (Fig. 1A). First, we engineered a human iPS cell line that constitutively expresses the mNG2_1-10_ fragment. We validated this cell line extensively, and named it “smNG2-P” (split mNeonGreen2 parental cell line). As a proof-of-concept, we efficiently targeted multiple genes for endogenous tagging with mNG2_11_, and clonally isolated several tagged iPS cell lines. To facilitate this process, we developed protocols for efficient single-cell isolation by FACS and screening of edited clones by Nanopore sequencing. Finally, we show how the endogenously tagged iPS cell lines can be used for live imaging studies of cytokinesis, which is a highly dynamic cellular process. Timelapse imaging of several weakly expressed cytokinesis genes that were endogenously tagged in iPSCs revealed new insights to how cytokinesis occurs in these cells. Imaging the more weakly expressed genes required the use of an image restoration algorithm to alleviate cell toxicity and obtain high quality images with high temporal resolution. This work provides the foundation for a genome-wide endogenous tag library in human stem cells and provides protocols to efficiently generate and study endogenous tags in human iPSCs.

**Figure 1.**
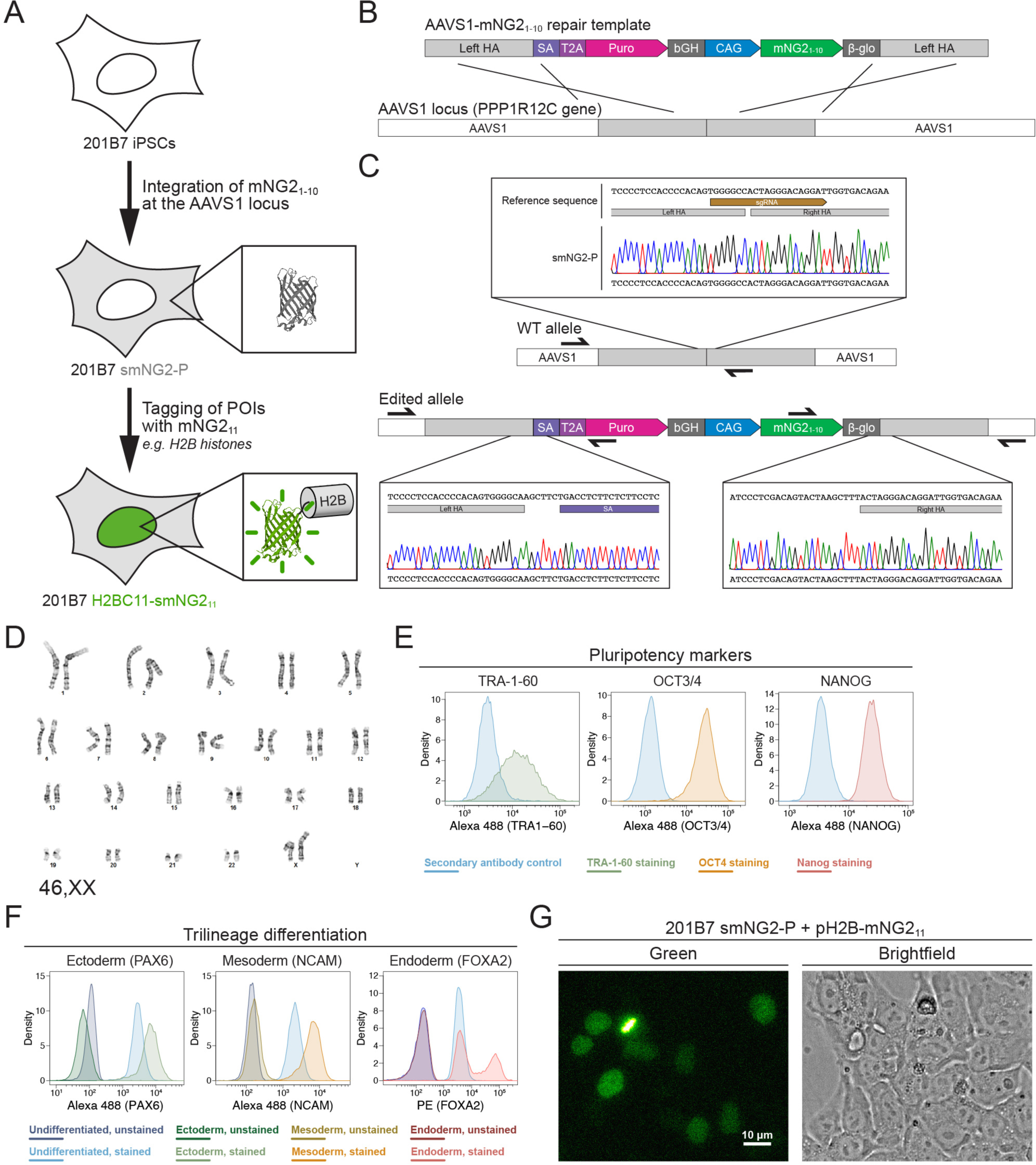
Generation of a split mNeonGreen human iPS cell line. **A)** A schematic representation of the split tagging strategy used in this study. An expression cassette carrying the mNG2_1-10_ fragment was integrated at the AAVS1 locus in 201B7 iPSCs to generate the parental split mNeonGreen2 cell line (smNG2-P). Proteins of interest were tagged endogenously with the mNG2_11_ fragment to visualize their expression and localization. The structure of mNeonGreen was generated from PDB (Protein Data Bank, structure identifier 5LTP; Clavel et al., 2016). **B)** A schematic of the mNG2_1-10_ expression cassette is shown with the CAG promoter (blue) and sequences for stable expression (dark grey), Puromycin resistance marker (pink), and homology arms (light grey) for integration at the AAVS1 locus. Abbreviations: SA = splicing acceptor, T2A = *Thosea asigna* virus 2A peptide, bGH = bovine growth hormone poly-adenylation signal, β-glo = rabbit β-globin poly-adenylation signal. **C)** Sequencing chromatograms show the edited AAVS1 allele junctions in the smNG2-P cell line (bottom) compared to the WT allele (top). **D)** Representative G-banding karyotype of the smNG2-P cell line shows that the edited cells have a normal karyotype (46, XX). **E)** Flow cytometry plots show smNG2-P cells stained for pluripotency markers TRA-1-60 (left, green), OCT3/4 (center, yellow) and NANOG (right, red) or a secondary antibody control (blue). **F)** Flow cytometry plots show differentiated smNG2-P cells stained for PAX6 (ectoderm; left, green), NCAM (mesoderm; center, yellow) and FOXA2 (endoderm; right, red) compared to undifferentiated cells (blue) and unstained controls (dark colors). **G)** Fluorescent and brightfield images show fluorescence complementation after transfecting smNG2-P cells with a plasmid expressing H2B fused to mNG2_11_. The scale bar is 10 microns.

## Results

### Generation of a split mNeonGreen iPS cell line for efficient endogenous tagging

Our first goal was to generate a human iPS cell line where mNG2_1-10_ is constitutively expressed. To do this, we generated a repair template that contains a mNG2_1-10_ expression cassette for integration into the AAVS1 locus, based on a previously published design (Fig. 1B; Oceguera-Yanez et al., 2016). This cassette contains the CAG promoter to drive high levels of mNG2_1-10_ expression, and resist silencing at the AAVS1 locus over time and during differentiation (Luo et al., 2014; Oceguera-Yanez et al., 2016). The cassette also includes the gene for Puromycin resistance, which is expressed upon integration at the AAVS1 locus via a splicing acceptor and self-cleaving T2A peptide (Fig. 1B). We introduced this mNG2_1-10_ expression cassette into the 201B7 cell line, which was generated by retroviral transduction of reprogramming factors into dermal fibroblasts and is well-characterized (Takahashi et al., 2007). After transfecting 201B7 human iPSCs with the AAVS1-mNG2_1-10_ repair template and an AAVS1-targeting Cas9/sgRNA RNP (ribonucleoprotein) complex, the cells were selected for Puromycin resistance and single-cell clones were isolated by FACS for screening. Clones were first screened by qPCR for the presence of the mNG2_1-10_ gene and the absence of the Ampicillin resistance (AmpR) gene, which was part of the backbone of the repair template (Fig. S1A-D). Clones that were positive for mNG2_1-10_ and negative for the AmpR gene were then screened for integration at the AAVS1 locus by junction PCR (Fig. S1E-H). We found 6 clones that were heterozygous for AAVS1-mNG2_1-10_ integration, and one clone was selected for full validation (clone 28).

### Validation of the smNG2-P iPS cell line

We further validated clone 28 to ensure that the cells are still healthy and carry only the desired AAVS1-mNG2_1-10_ integration. Sequencing of the AAVS1 locus revealed that one allele contains the mNG2_1-10_ expression cassette, while the other allele is unedited (Fig. 1C). To ensure that only one copy of the cassette was integrated properly, we measured the genomic copy number for mNG2_1-10_ and for the AmpR gene by digital PCR (dPCR). Our data confirmed the presence of a single copy of mNG2_1-10_ with no ectopic integration (Fig. S2A). Next, 12 sites predicted to be susceptible to off-target editing based on previous studies and prediction software were selected for sequencing (Concordet & Haeussler, 2018; Hsu et al., 2013; Wang et al., 2014). We found no differences in the sequence of the 12 off-target sites between the WT 201B7 cell line and our edited cells (Fig. S2B-M). To ensure that the clone 28 cells have a normal genome, we performed G-banding karyotyping and found no chromosomal abnormalities (46, XX karyotype; Fig. 1D). To verify that clone 28 cells are undifferentiated, we used immunofluorescence staining and flow cytometry with antibodies specific for TRA-1-60, OCT3/4 and NANOG (Fig. 1E). We found that all three pluripotency markers were expressed. We also found that clone 28 cells retain pluripotency by successfully differentiating them into ectoderm, mesoderm and endoderm, as determined by immunofluorescence staining and flow cytometry with antibodies specific for PAX6 (ectoderm), NCAM (mesoderm) and FOXA2 (endoderm; Fig. 1F). Clone 28 cells also have normal iPSC morphology and tested negative for mycoplasma contamination (Fig. S2N-O). Finally, we verified that mNeonGreen fluorescence could be reconstituted in the presence of the mNG_11_ fragment. Transfection of clone 28 cells with a plasmid expressing H2B (histone) fused to the mNG_11_ fragment resulted in the expression of a fluorescent signal localized to chromatin as expected for the functional complementation of the mNG2_1-10_ and mNG2_11_ fragments (Fig. 1G). After these validation and quality control steps, we chose clone 28 as the AAVS1-mNG2_1-10_ cell line, which we hereafter refer to as “smNG2-P” (split mNeonGreen2 parental cell line).

### Efficient endogenous tagging with mNG2_11_ in smNG2-P cells

Since fluorescence could be reconstituted by expressing the mNG_11_ fragment in the smNG2-P cell line, we aimed to integrate mNG2_11_ into different endogenous loci. We selected 17 genes for tagging, some of which had been previously endogenously tagged with the split mNG system (*e.g.* ACTB, TUBA1B, KRT18; Cho et al., 2022) or with full-length fluorescent proteins (*e.g.* H2BC11, ACTB, ANLN, RHOA; Husser et al., 2022; Roberts et al., 2017), while other genes had not been tagged previously (*e.g.* RACGAP1, KRT5, TUBB3, FOXA2). We also expected some of these genes to be expressed at varying levels in iPSCs. The expression levels for these genes are shown on a graph of RNA-seq data from (Iwasaki et al. 2022 Fig. S3A). While ACTB and TUBA1B should be highly expressed, there is a shift to lower expression levels for KRT18, RHOA and H2BC11, while SOX2, CDH1, NES, TUBB3, ANLN and RACGAP1 are all weakly expressed. Finally, NCAM1, FOXA2, PAX6, TBXT, KRT5 and KRT14 should be silent in iPSCs as they are only expressed in other cell types (Fig. S3A). Altogether, these genes cover a range of expression levels, localization patterns and involvement in different cellular processes. For each protein, we selected the N- or C-terminus for tagging based on published studies, and we used a short flexible linker (GGG; Feng et al., 2017; Kamiyama et al., 2016; Leonetti et al., 2016) to minimize disruption of the tagged protein. We designed ssODNs to introduce the mNG2_11_ tag at these loci, and transfected them into smNG2-P cells as Cas9/sgRNA RNP complexes. Eight days after transfection, we quantified the proportion of WT, indel and tagged alleles in the edited cell populations by Nanopore sequencing (Fig. 2A-B). We found that Cas9 targeting was efficient for all loci, as shown by the high proportion of indels in the edited cell populations. However, the proportion of tagged alleles was variable, with 37.5% of CDH1 alleles tagged, while only 0.69% of TUBA1B alleles were tagged (Fig. 2B). This data shows that while endogenous tagging with mNG2_11_ can be highly efficient, it varies considerably for each locus.

**Figure 2.**
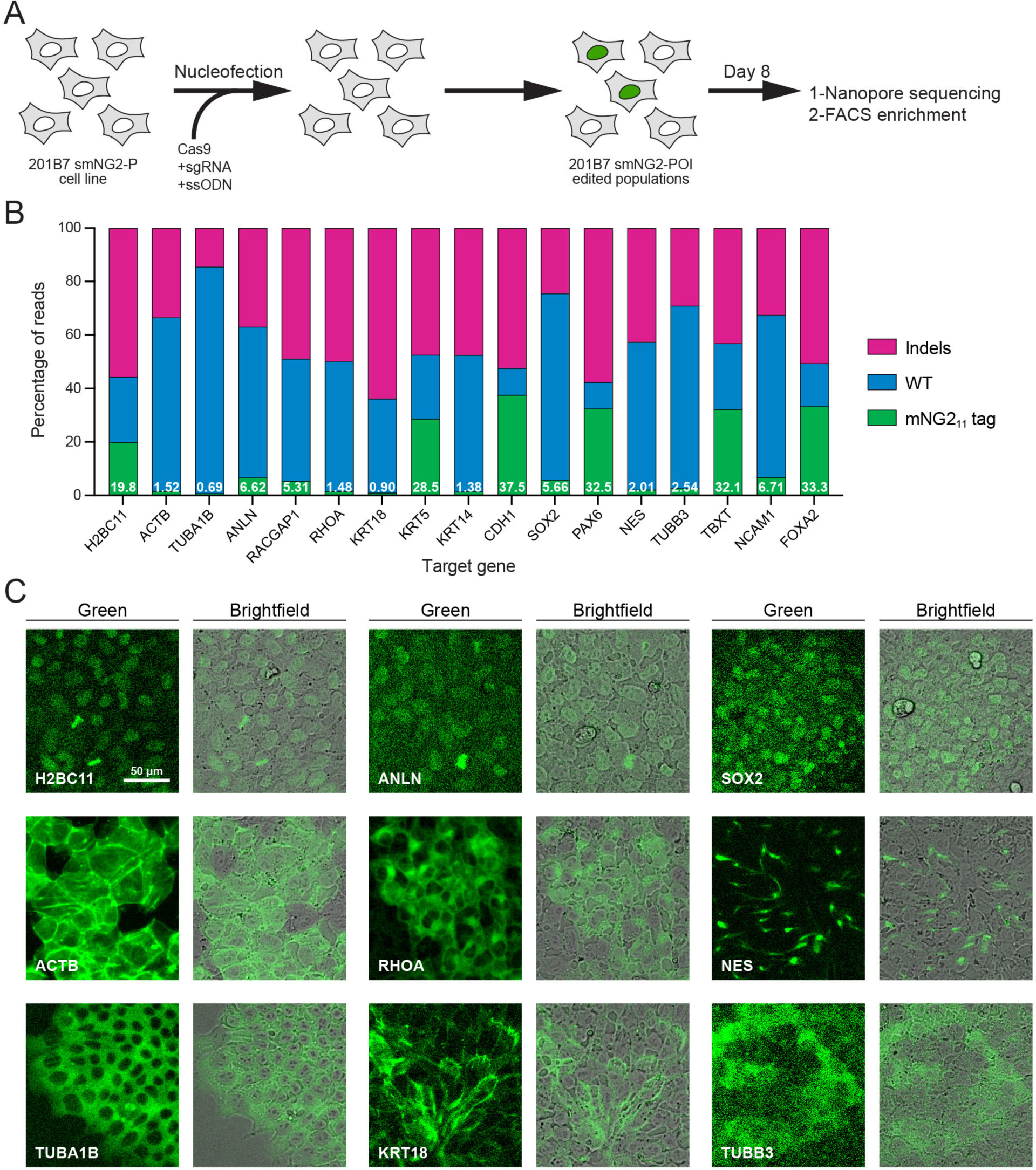
Efficient endogenous tagging with mNG2_11_ in smNG2-P cells. **A)** A schematic shows the workflow used to tag proteins of interest with the mNG2_11_ fragment. smNG2-P cells were transfected with Cas9/sgRNA RNPs and a ssODN repair template to integrate mNG2_11_ at a target locus. After recovery, the edited populations were frozen, then assessed by Nanopore sequencing and flow cytometry. **B)** A bar graph shows the distribution of alleles (WT, indels or mNG2_11_ tagged) in edited populations. The percentage of mNG2_11_ alleles is indicated in white for each gene. **C)** Fluorescent and brightfield images show populations of tagged cells after enrichment by FACS as indicated. The scale bar is 50 microns.

Next, we monitored the reconstitution of mNeonGreen fluorescence in edited populations by flow cytometry and fluorescence microscopy. As expected for genes that are expressed in iPSCs, we observed a fluorescent signal in H2BC11, ACTB, TUBA1B, ANLN, RHOA, KRT18, SOX2, NES and TUBB3-tagged populations (Fig. S3B-T). Tagged cell populations were enriched by FACS, and microscopy was used to determine if their localization was consistent with prior studies (Fig. 2C). As expected, the fluorescent signal from H2B histone was nuclear (Viana et al., 2023), β-actin formed filaments and was enriched at the cortex (Roberts et al., 2017; Viana et al., 2023), α-tubulin was cytosolic and enriched in filaments and mitotic spindles (Roberts et al., 2017; Viana et al., 2023), anillin was nuclear and enriched in the furrow of dividing cells (Hesse et al., 2012; Husser et al., 2022), RhoA was cytosolic and weakly enriched at the cortex (Husser et al., 2022; Yonemura et al., 2004), keratin 18 formed cortical filaments (Maurer et al., 2008), SOX2 was nuclear (Allencell.org; Strebinger et al., 2019), Nestin formed distinct filaments (Kuang et al., 2019), and β-3-tubulin was weakly expressed and cytosolic (Guo et al., 2010; Turac et al., 2013). For these proteins, the mean fluorescence intensity of tagged cells correlated well with their expected relative expression levels (Fig. S3A and U). Despite the high tagging efficiency observed for CDH1, we did not observe any fluorescent signal by flow cytometry (Fig. S3K). However, microscopy revealed that E-cadherin-mNG2_11_ was enriched at cell junctions (Fig. S3V; Allencell.org; Aban et al., 2021; Cumin et al., 2022). This suggests that weakly expressed proteins may require different methods to verify expression following endogenous tagging. However, we did not observe any fluorescent signal from tagged Cyk4 (RACGAP1 gene) by flow cytometry or microscopy (Fig. S3G). The lack of signal could be due to the low tagging efficiency (Fig. 2B), and/or the weak expression of Cyk4 in stem cells (Fig. S3A; Iwasaki et al., 2022). There was no fluorescent signal in the KRT5, KRT14, PAX6, TBXT, NCAM1 and FOXA2-edited populations, as expected for differentiation markers (Fig. 2B and S3). We observed nuclear fluorescence by microscopy in FOXA2-tagged cells after induction into endoderm (Fig. S3W-X), but we did not investigate the expression of the others as this goes beyond the scope of our study.

### Efficient clonal isolation and screening of tagged iPS cell lines

Next, we isolated 5 clonal cell lines expressing H2B histones, β-actin, α-tubulin, anillin and RhoA tagged with mNG2_11_, as these proteins have distinct localization patterns during cell division (Beaudet et al., 2017; Husser et al., 2022; Rodrigues et al., 2015; van Oostende Triplet et al., 2014). We optimized a protocol to efficiently recover single-cell clones after FACS (shown in Fig. 3A), resulting in an average recovery of 61.4% in 96-well plates across the 5 genes (Fig. S4A). To facilitate screening of clonal recovery in 96-well plates, we developed an ImageJ macro that automatically identifies positive and negative wells from 96-well plate images (Fig. S4B-G). Isolated clones were then screened based on their genotype by multiplexed Nanopore sequencing. We found that clones had diverse genotypes with alleles containing mNG2_11_, mutated mNG2_11_, indel mutations and WT sequences (Fig. 3B). The diversity of genotypes amongst tagged cells suggests that clonal isolation is important to obtain a high-quality homogeneous population of cells for further study. We obtained both homozygous and heterozygous clones carrying smNG2-anillin and smNG2-RhoA, and we found homozygous clones that were brighter than heterozygous ones (Fig. S4H-I). This suggests that the complementation between mNG2_1-10_ and mNG2_11_ occurs efficiently over a range of mNG2_11_ expression, since anillin is expressed weakly and RhoA is expressed more strongly in iPSCs. We also observed some homozygous clones that were not brighter than the corresponding heterozygous clones, which could be due to undetected by- products of CRISPR or clonal variation in protein expression. This variability underlines the importance of screening and validating edited clones. For each tagged protein, a single clone was selected for further study (Fig. 3C-G). These cell lines were also tested for mycoplasma contamination (Fig. S4J). Fluorescent images of the final mNG2_11_-tagged cell lines acquired using confocal sweptfield microscopy are shown in Figure 3H. Consistent with the expected localization for these proteins, H2B was nuclear during interphase and localized to condensed chromatin during mitosis, actin was cortical in both interphase and mitotic cells, tubulin was cytosolic during interphase and localized to mitotic spindles, anillin was nuclear during interphase and enriched in the cleavage furrow during cytokinesis, and RhoA was cytosolic during interphase and cortically enriched during mitosis.

**Figure 3.**
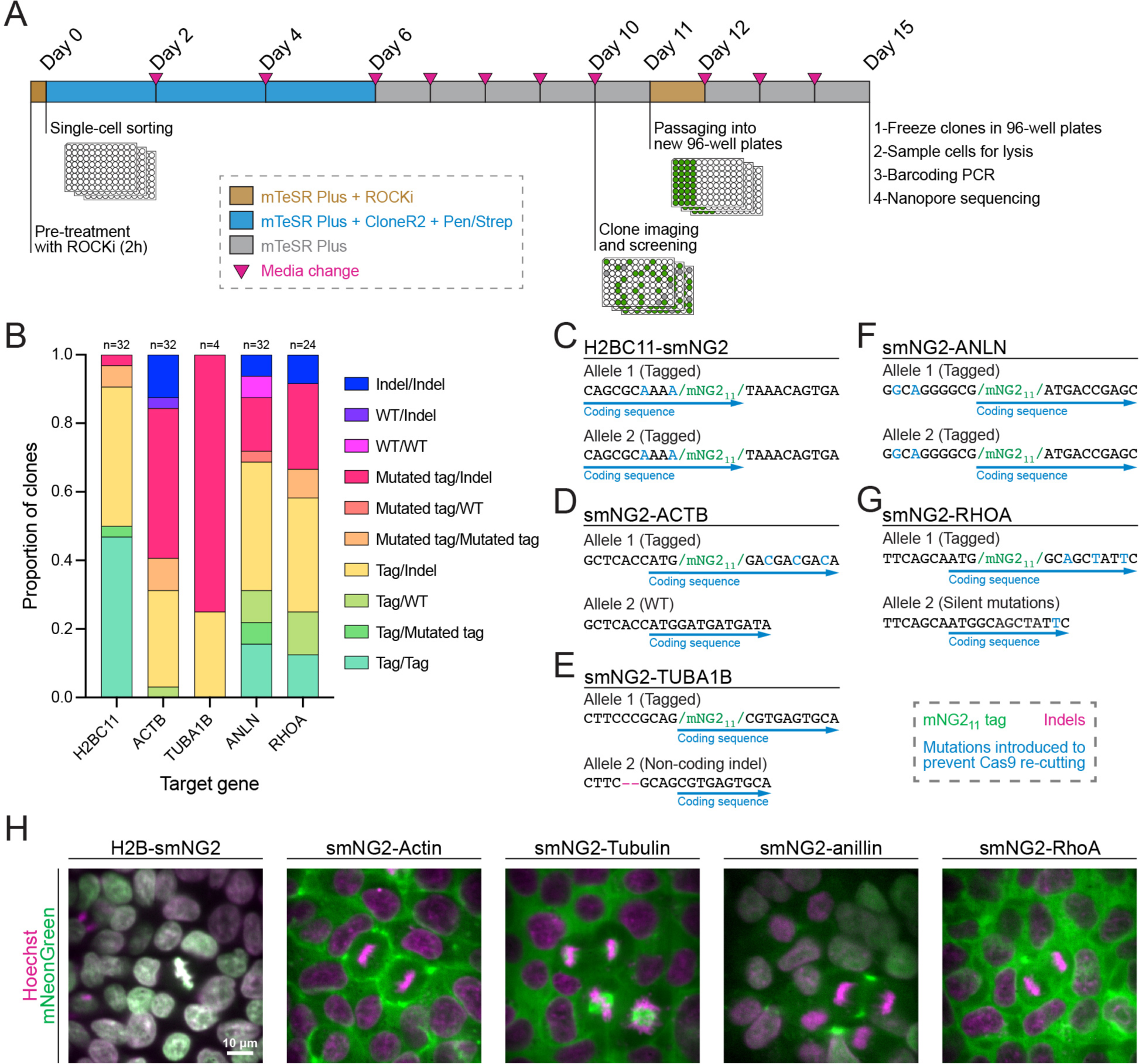
Efficient clonal isolation and screening of tagged iPS cell lines. **A)** The protocol used for single-cell isolation and the recovery of tagged iPSC clones is shown. Tagged cell populations were pre-treated with ROCK inhibitor Y-27632 for 2 hours before sorting individual cells into 96-well plates by FACS. Cells were kept in recovery media for 6 days, then colonies were screened on day 10 to select those for further passaging into 96-well plates on day 11. Fully-grown clones were frozen on day 15, and cells were genotyped by barcoding PCR followed by Nanopore sequencing. **B)** A stacked bar graph shows the distribution of genotypes inferred from multiplexed Nanopore sequencing for isolated clones of tagged H2BC11, ACTB, TUBA1B, ANLN and RHOA (sample size is indicated above each bar). **C-G)** The genotypes of the final tagged cell lines are shown: H2BC11-smNG2 (C), smNG2-ACTB (D), smNG2-TUBA1B (E), smNG2-ANLN (F) and smNG2-RHOA (G). The target gene coding sequence is in blue, indel mutations are in red and mNG_11_ tag in green. **H)** Fluorescent images show the localization of smNG2 (green) and DNA (stained with Hoechst; magenta) in the final tagged cell lines. The scale bar is 10 microns.

### Image reconstitution for live imaging of cellular processes in iPSCs

The endogenously tagged iPS cell lines can be used to study the mechanisms controlling different cellular processes in healthy cells and how they vary with cell type. Our knowledge of human cytokinesis is derived from transformed and/or cancerous, differentiated cell lines and has not been studied in human pluripotent stem cells. Cytokinesis is dynamic and requires imaging over long periods of time (tens to hundreds of minutes) at frequent intervals. To study cytokinesis in human iPSCs, we imaged the mNG2_11_-tagged cell lines from metaphase onwards. After a few minutes of imaging using standard optical settings, iPSCs stopped dividing and detached from the coverslip, independently of laser wavelength (data not shown), suggesting that iPSCs are more sensitive to photo-toxicity than transformed and cancerous cell lines. Moreover, endogenous protein levels are low, requiring higher exposure times. To overcome this issue, we optimized the settings to reduce exposure time and laser power, decrease z-stack depth and increase imaging intervals so that cells could be imaged for more than 80 minutes. However, while we could detect sufficient levels of actin using these imaging conditions, proteins that are more weakly expressed were more challenging to detect (Fig. 4A). Indeed, the signal-to-noise ratio was low for H2B, α-tubulin, anillin and RhoA in images acquired with low-exposure settings that supported cell survival, compared to images acquired using high-exposure settings that did not support cell viability (Fig. 4A). To overcome this issue, we used deep learning-based image restoration to obtain high resolution images from those with low signal-to-noise ratios caused by low-exposure settings. We trained a CARE (Content-Aware image REstoration) neural network on sets of matched high- and low-exposure images for each cell line (Fig. 4B; Weigert et al., 2018), and used the trained model to restore timelapse movies acquired with low-exposure settings (Fig. 4C-G). The signal-to-noise ratio was drastically improved for timelapse images of H2B, α-tubulin, anillin and RhoA (Fig. 4C-G). Most importantly, this approach allowed us to image live iPSCs with high temporal resolution.

**Figure 4.**
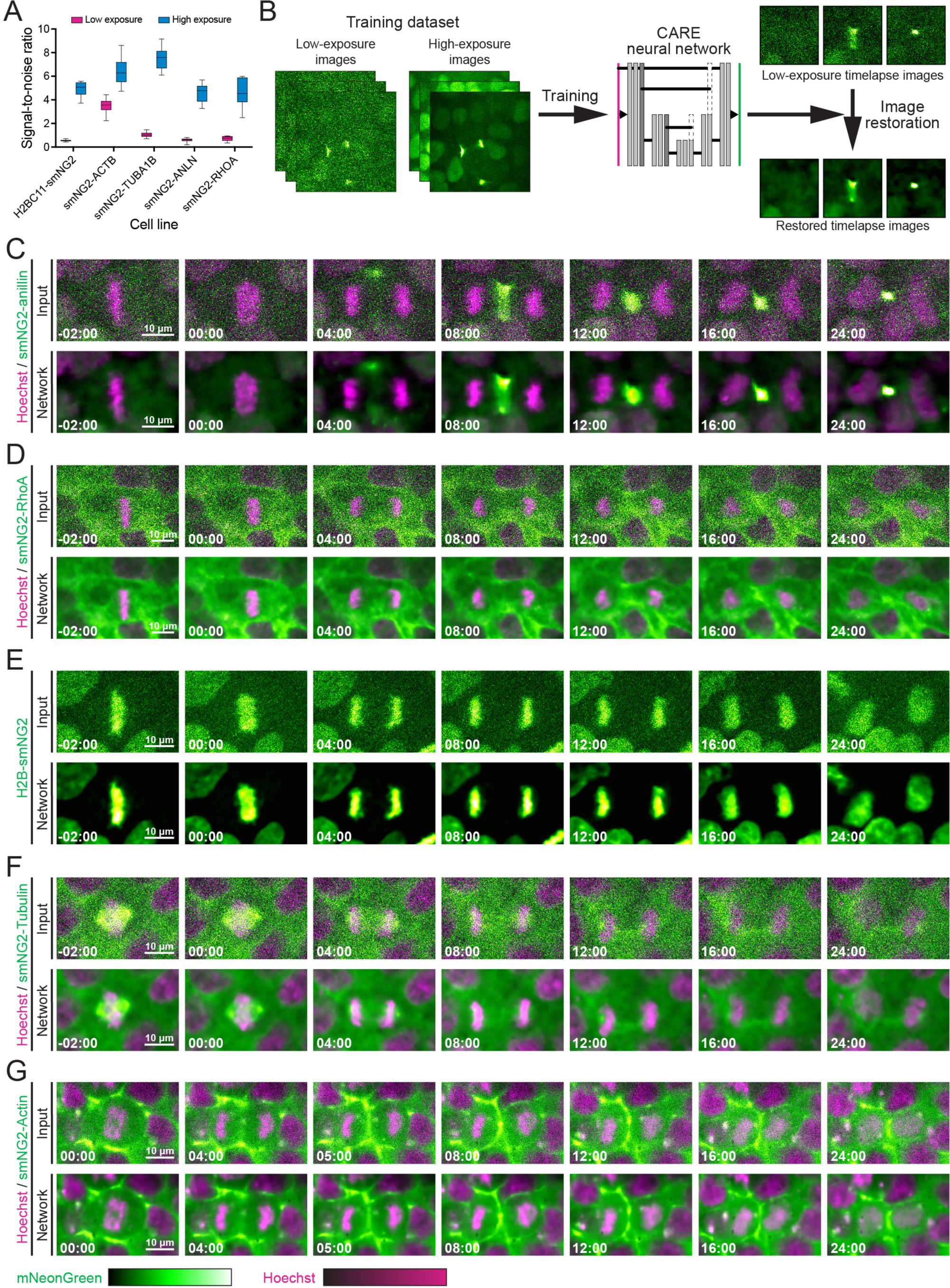
Image restoration for live imaging and quantitative measurements in iPSCs. **A)** A graph shows the signal-to-noise ratio measured from low-(pink) and high-exposure (blue) images of the tagged iPS cell lines as indicated. **B)** A schematic shows the training and application of the CARE neural network for image restoration. The neural network developed by Weigert et al. (2018) was trained on sets of low- and high-exposure images for each tagged cell line and used to restore timelapse movies. **C-G)** Comparisons of raw (Input; top panel) and restored timelapse images (Network; bottom panel) by the CARE neural network are shown for iPSCs expressing smNG2-anillin (C), smNG2-RhoA (D), H2B-smNG2 (E), smNG2-Tubulin (F) and smNG2-actin (G) undergoing cytokinesis (smNG2 in green; DNA stained with Hoechst in magenta). The scale bars are 10 microns, and time is relative to anaphase onset (00:00).

With the ability to image key cytoskeletal and cytokinesis regulators such as RhoA and anillin, we characterized cytokinesis in iPSCs for the first time. In metazoans, cytokinesis initiates by the assembly of a RhoA-dependent contractile ring in anaphase which ingresses during telophase to pinch in the membrane (Fig. 5A), and then transitions to a stable midbody for abscission (Ozugergin & Piekny, 2022; Pollard & O’Shaughnessy, 2019). Anillin crosslinks key components of the ring, membrane and spindle for ring positioning, ingression and midbody formation (Green et al., 2012; Piekny & Maddox, 2010). We recently found that ring assembly kinetics and ingression varies among a few transformed and/or cancerous human cell types, and correlates with the localization of anillin (Husser et al., 2022). Characterizing cytokinesis in iPSCs will be an important starting point to build new knowledge of how cytokinesis is regulated in healthy, non-transformed cells. First we measured the duration of contractile ring ingression in anillin-tagged iPSCs (18.6 ± 4.0 minutes; Fig. 5B). We then measured the localization of anillin, RhoA and actin in metaphase and throughout cytokinesis. In metaphase cells, we found that anillin is cytosolic, while RhoA is weakly enriched at the cortex and actin is strongly enriched at the cortex (Fig. 4C, D, G, and 5C). During mitotic exit, anillin accumulates at the equatorial cortex ∼4 minutes after anaphase onset, while RhoA accumulates by ∼6 minutes, and actin remains strongly cortical with some equatorial enrichment by ∼4-5 minutes (Fig. 4C, D, G and 5D). We also found that actin is enriched at cell junctions, which remodel between adjacent cells during cytokinesis (Fig. S5A). In an extreme example, junctions disappeared, and foci appeared adjacent to the site of ingression, likely resulting from the mechanical response to the forces generated by the furrow (Fig. S5A). We then measured the breadth of cortically enriched anillin and RhoA at the onset of ingression and found that anillin forms a narrow peak (4.1 ± 0.7 μm), while RhoA localizes more broadly (6.6 ± 2.2 μm; Fig. 5E-F). The same result was obtained when measuring furrow breadth as a percentage of cortex length, showing that the difference between RhoA and anillin localization is independent of cell size (Fig. S5B). Since RhoA and anillin are expected to closely co-localize (Piekny & Maddox, 2010), we investigated this discrepancy by looking at the central spindle and astral microtubules in high-exposure images of tagged tubulin cells (Fig. 5G). We found that cells have a poorly-organized central spindle in anaphase (Fig. 5G-H), and the astral microtubules appear to contact the equatorial cortex (Fig. 5G). This data reveals new differences in how cytokinesis occurs in iPSCs and demonstrates the utility of endogenous tags to study cellular processes by live imaging, which also can be done in differentiated cell types with isogenic backgrounds.

**Figure 5.**
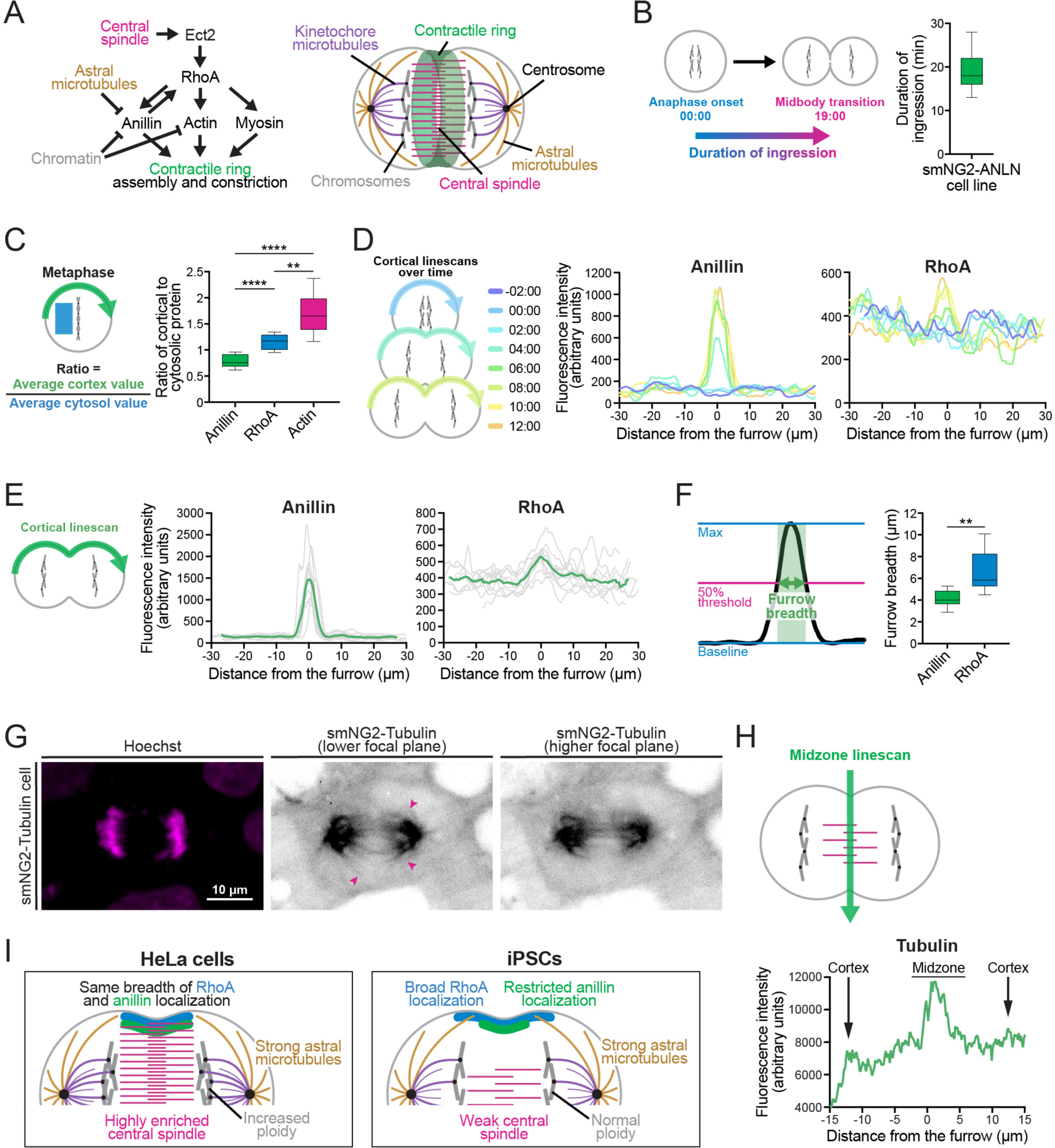
Cytokinesis is uniquely regulated in iPSCs. **A)** Schematics show the core pathways regulating contractile ring assembly and ingression (left) and their localization inside a late anaphase cell (right). **B)** A schematic (left) shows how the duration of ingression was measured in anillin-tagged iPSCs (right; n = 39). **C)** A schematic (left) shows how the ratio of cortical to cytosolic protein was measured in metaphase cells for smNG2-anillin, smNG2-RhoA and smNG2-actin (right; n = 10 for each line). **D)** A schematic (left) shows the location and timing of the linescans used to plot the fluorescence intensity along the cortex in single iPSCs for smNG2-anillin (center) and smNG2-RhoA (right). Timepoints are shown in different colors as indicated in the scale. **E)** A schematic (left) shows the location of the linescans used to plot the intensity of fluorescence along the cortex at the onset of furrowing for smNG2-anillin (center) and smNG2-RhoA (right; n = 10 for each). **F)** A schematic (left) shows how the breadth of protein localization at the equatorial cortex at furrow initiation was calculated for smNG2-anillin and smNG2-RhoA (n = 10 for each). **G)** Fluorescent images show the location of DNA (left; stained with Hoechst) and smNG2-Tubulin (grayscale) in a cell. For tubulin, a lower focal plane (center) is shown to highlight astral microtubules (indicated by pink arrows), and a higher focal plane (right) is shown to highlight the central spindle. The scale bar is 10 microns. **H)** A schematic (top) shows the location of the linescan used to plot the fluorescence intensity along the midzone of a cell expressing smNG2-tubulin in anaphase (bottom). **I)** A schematic shows a comparison of conventional view of cytokinesis (HeLa cell; left) and an iPS cell (iPSCs; right). HeLa cells have a strongly enriched central spindle, which keeps the equatorial of active RhoA narrow, and astral microtubules that extend all the way to the cortex and restrict the localization of anillin to the same zone as RhoA. iPSCs have a weakly enriched central spindle, resulting in a broader zone of active RhoA, and prominent astral microtubules that keep anillin in a narrow equatorial zone. Box plots in B, C and F show the median line, quartile box edges and minimum and maximum value whiskers. Statistical significance was determined by Brown-Forsythe and Welch’s ANOVA for C and Welch’s t test for F (ns, not significant; * p≤0.05; ** p≤0.01; *** p≤0.001; **** p≤0.0001).

## Discussion

Recent advances in genetic engineering have accelerated the development of tools for fundamental research. Endogenous tags are particularly valuable as they enable the visualization of protein behavior in live cells at endogenous expression levels. Self-complementing split fluorescent proteins have been used for efficient endogenous tagging, and this approach was recently used for the construction of the first library of endogenous tags in HEK293 cells, with 1,310 proteins tagged in mixed populations enriched by FACS (Cho et al., 2022). While this library is a powerful resource for the community, researchers will need to generate single clones of individual tagged proteins before they can be studied. Further, the knowledge generated by this library is restricted to this single cell type. Given the predicted diversity in mechanisms controlling cellular processes across cell types, there is a need to also generate endogenous tags in stem cells that can be differentiated into multiple cell types. In this study, we generated a fully validated human iPS cell line expressing mNG2_1-10_ for efficient endogenous tagging with mNG2_11_ in human stem cells capable of differentiating into any cell type. The 201B7 smNG2-P cell line is heterozygous for the mNG2_1-10_ expression cassette which was integrated at the AAVS1 safe-harbor locus. Compared to lentiviral delivery, this design minimizes the risk of mNG2_1-10_ silencing and provides high and consistent expression of mNG2_1-10_ across all cells in the population, even over time and during differentiation (Oceguera-Yanez et al., 2016). We tagged multiple genes with mNG2_11_, with efficiencies ranging from 0.69% to 37.5% measured by Nanopore sequencing (Fig. 2B). Tagging efficiency with mNG2_11_ in iPSCs was lower than in mouse embryos (40 to 100% of injected embryos; O’Hagan et al., 2021), and lower than tagging with GFP_11_ in HEK293 cells (<1% to 56% tagged alleles; Cho et al., 2022). However, endogenous tagging with mNG2_11_ was overall more efficient than endogenous tagging with full-length mEGFP in iPSCs (mostly <0.1 to 4%, and up to 24% GFP-positive cells; Roberts et al., 2017). Although it is difficult to compare editing efficiencies across studies, endogenous tagging by HDR is very inefficient. Our results are consistent with iPSCs being more challenging to edit compared to other cell types, and mNG2_11_ integrating at higher frequency than larger tags. The split mNeonGreen system also enables endogenous tagging on the scale of hundreds to thousands of proteins in parallel. For comparison, endogenous tagging with full-length fluorescent proteins requires the assembly of large repair templates, which limits the number of proteins that can be tagged at once. We also found that low endogenous expression levels can limit the detection of tagged proteins by flow cytometry, similar to previous studies (Cho et al., 2022; Leonetti et al., 2016; O’Hagan et al., 2021), requiring microscopy with highly sensitive cameras or detectors. Alternative methods such as fixation and antibody staining can also be used for more sensitive detection of weakly expressed endogenous tags (O’Hagan et al., 2021). Importantly, the expression levels of 10 mNG2_11_-tagged proteins in iPSCs (Fig. 2C and S3U) were consistent with RNA-seq data (Fig. S3A; Iwasaki et al., 2022). Since these proteins showed unique localizations patterns consistent with their function, it is unlikely that the mNG2_11_ tag affected their function. Interestingly, Nestin and β-3-tubulin were expressed at low levels in undifferentiated iPSCs, despite being commonly used as ectodermal and neuronal differentiation markers. This result is consistent with a previous report of their expression in some iPS cell lines (Kuang et al., 2019) and with published RNA-seq and proteomic data for the 201B7 cell line (Iwasaki et al., 2022). The detection of a broader range of endogenous protein expression in live cells will require further improvements in the brightness of fluorescent proteins or in the sensitivity of detectors.

We also found that edited cells had diverse genotypes caused by gene editing, including alleles carrying mutated mNG2_11_ tags and indel mutations, warranting clonal isolation. Such mutations have been reported previously but are not fully understood (Burgio & Teboul, 2020; Skryabin et al., 2020). Single-nucleotide mutations may come from mutations introduced during ssODN synthesis. Meanwhile, ssODN re-arrangements could be caused by microhomologies, repair by NHEJ instead of HDR, or a combination of NHEJ and HDR repair (Skryabin et al., 2020). Unwanted editing outcomes are ignored when looking at pooled populations, and we recommend isolating and screening clonal cell lines to perform quantitative studies of desired cellular processes, as they provide a more reliable readout of protein expression, where variations in signal between cells within a population represent true cell-to-cell variability. Clonal isolation is notoriously inefficient in iPSCs, which are programmed to undergo dissociation-induced apoptosis upon loss of attachment or cell-cell contact (Bhargava et al., 2022; Chen & Pruett-Miller, 2018; Singh, 2019; Tristan et al., 2023; Watanabe et al., 2007). We optimized a protocol for efficient single-cell recovery of 201B7 iPSCs after FACS, with up to 80% of wells in a 96-well plate showing clonal growth after 10 days (Fig. 3A and S4A). We also created a macro for image-based colony screening in 96-well plates (Fig. S4B-G), and used multiplexed Nanopore sequencing for high-throughput screening of selected colonies, as many amplicons can be sequenced simultaneously (Whitford et al., 2022). Automated instrumentation could be used to further increase the scale of clonal cell isolation, and to isolate cell lines where transcriptionally silent genes are tagged endogenously (Roberts et al., 2019). After clonal isolation, we found that the cells in each edited line had consistent, stable fluorescence. Further, we observed different levels of fluorescence across a range of protein expression levels (Fig. S3A-T), showing that mNG2_1-10_/mNG2_11_ complementation is comparable to full-length fluorescent proteins. Importantly, the stability of fluorescence levels in endogenously tagged cell lines and the absence of localization artefacts compared to over-expressed transgenes makes this system attractive for quantitative measurements in live cells (Husser et al., 2022).

The goal of endogenous tagging in iPSCs is to enable live imaging studies of diverse cellular processes for comparative studies among different human cell types. We found that iPSCs are particularly sensitive to phototoxicity, making it difficult to image weakly expressed endogenous tags over longer periods of time or with short time intervals. This sensitivity has not been previously reported in human or in mouse stem cells, likely because imaging conditions are rarely reported and vary with different setups. In addition, timelapse imaging often involves the use of over-expressing transgenes or dyes, which generate higher fluorescent signal intensity and enable the use of optical settings with lower laser power and exposure time (Chaigne et al., 2021; Hesse et al., 2012; Roberts et al., 2017). Since most proteins are expressed weakly in the endogenous context, improvements in fluorophore brightness, imaging conditions or image processing are required. Since many labs have access to spinning disk confocal or epifluorescence widefield microscopes, image restoration methods provide a way to obtain high-quality images of iPSCs without compromising on temporal resolution. We used CARE, a content-aware image restoration neural network developed by Weigert et al. (2018), where training datasets are created with matched images obtained using low- and high-exposures (Fig. 4). With this network, cells can be imaged over extended periods of time with high temporal resolution using low exposure settings, and image files can be restored to generate high-quality movies.

As a proof-of-concept, we characterized cytokinesis in live human iPSCs for the first time. Most of our knowledge of human cell cytokinesis was obtained from studies using HeLa cells, yet the mechanisms regulating are expected to vary with cell type (Husser et al., 2022; Ozugergin & Piekny, 2022). As a result, we do not have detailed knowledge of how cytokinesis occurs in most healthy human cell types. CARE allowed us to measure the timing of cytokinesis and the localization of core cell components (actin, tubulin) and cytokinesis proteins (RhoA, anillin) in live human iPSCs (Fig. 5). We found that anillin-tagged iPSCs completed ingression in 18.6 ± 4.0 minutes; faster than HepG2 and HEK293 cells, slower than HCT116 cells, and similar to HeLa cells (Fig. 5B; Husser et al., 2022). Anillin is recruited to a narrow band at the equatorial cortex ∼4 minutes after anaphase onset, while RhoA is also enriched but only after ∼6 minutes and is more broadly distributed in iPSCs, while in HeLa cells they are both enriched after ∼6 minutes and have the same breadth (Fig. 5D-E; (Husser et al., 2022; Piekny & Maddox, 2010). Active RhoA is generated at the equatorial cortex by the guanine nucleotide exchange factor Ect2 which is activated by Cyk4 at the central spindle (Koh et al., 2022; Mahlandt et al., 2021; Yuce et al., 2005). In HeLa cells, the central spindle starts to form immediately after anaphase onset and reaches the cortex by late anaphase (∼6 minutes) (van Oostende Triplet et al., 2014). Thus, the weak central spindle in anaphase iPSCs may explain the broad localization of RhoA (Fig. 5G-H). In support of this, perturbations that weaken the central spindle in HeLa cells result in more diffuse Ect2-Cyk4 complexes and a broader zone of active RhoA (Adriaans et al., 2019; Kotynkova et al., 2016; Su et al., 2011; Yuce et al., 2005). Active RhoA is also required for the cortical recruitment of anillin, yet the localization of anillin in iPSCs is more narrow compared to the other human cell lines that we previously characterized (Fig. 5F; Husser et al., 2022; Piekny & Glotzer, 2008). In iPSCs, the astral microtubules extended to the cortex in the equatorial region, which could restrict anillin localization. In HeLa cells, we previously showed that perturbations that cause an increase in astral microtubules decreases the breadth of anillin (van Oostende Triplet et al., 2014). However, interactions with other proteins or phospholipids could further restrict the localization of anillin in iPSCs (Ozugergin & Piekny, 2022). Previously, we speculated that the breadth of anillin localization inversely correlates with the speed of ingression. The iPSCs seem to differ from this model, suggesting that other factors should also be considered, such as the distribution of other actin crosslinkers or cortical flows (Khaliullin et al., 2018; Leite et al., 2020; Osorio et al., 2019; Reymann et al., 2016; Sobral et al., 2021; Spira et al., 2017). Thus our findings reveal how cytokinesis occurs in a non-transformed human stem cell, and provide a framework to reveal how the mechanisms controlling cytokinesis vary with different cell types in an isogenic context.

Our work combines approaches to alleviate some of the challenges of gene editing in human stem cells. High transfection and editing efficiencies can be achieved in iPSCs by using electroporation and delivering Cas9/sgRNA RNP complexes instead of plasmids (Kim et al., 2014; Liang et al., 2015). Using a short tag that can be carried on single-stranded repair templates (ssODNs) bypasses the requirement for cloning dsODNs with large homology arms. Tagging efficiency is further increased by using NHEJ inhibitors (Chu et al., 2015; Maruyama et al., 2015; Maurissen & Woltjen, 2020; Schimmel et al., 2023). Cells with mutated mNG2_11_ tags can be eliminated by screening single-cell clones based on genotype, using single cell isolation and image-based colony screening in 96-well plates, combined with multiplexed genotyping of clones by Nanopore sequencing. Finally, we showed how iPSCs could be imaged with high temporal resolution using CARE for studies of cytokinesis, an essential dynamic cellular process. Altogether, this work provides the basis for a high-quality endogenous tag library in human iPSCs, which will be used to study protein function in human stem cells and across human cell types *in vitro*.

## Materials and Methods

### Cell culture

201B7 iPSCs were obtained from ATCC. Cells were cultured in 6-well plates coated with iMatrix-511 Silk (Nippi) as per manufacturer’s protocol, in mTeSR Plus media (STEMCELL Technologies). The cells were kept at 37°C and 5% CO_2_. The cells were passaged every 3-4 days by dissociating them using Accutase (STEMCELL Technologies) according to manufacturer’s protocol. After dissociation, the cells were resuspended and 50,000-100,000 cells were transferred into fresh matrix-coated plates with mTeSR Plus media containing 10 μM Y-27632 (ROCK inhibitor, STEMCELL Technologies). Fresh media without ROCK inhibitor was fed the next, and the media was replaced daily until the next passage.

### Constructs

All plasmids were maintained and cloned in *Escherichia coli* DH5α unless specified otherwise. Point mutations were introduced into pNCS-mNeonGreen (Allele Biotechnology) by PCR-based site-directed mutagenesis to convert it to mNeonGreen2 (Feng et al., 2017). The mNG2_1-10_ fragment was then amplified by PCR. The left and right homology arms for the AAVS1 locus and the puromycin gene were amplified individually from the pAAVS1-P-CAG-GFP plasmid (Addgene #80491; Oceguera-Yanez et al., 2016). These fragments were cloned into the pYTK089 backbone by Golden Gate using a standard protocol (Lee et al., 2015). Since the CAG promoter was recalcitrant to amplification by PCR, it was digested directly from pAAVS1-P-CAG-GFP using BclI and EcoRI, gel extracted, and ligated into the pre-digested promoter-less plasmid to generate the pAAVS1-P-CAG-mNG2_1-10_ repair template (Addgene #206042). For BclI digestion, the plasmids were extracted from methylase-deficient *E. coli* ER2925. pH2B-mNG2_11_ (Addgene #206043) was generated by amplifying the H2B gene from an H2B-mRuby plasmid (Beaudet et al., 2017) with primers designed to add mNG2_11_ on the end of H2B. This gene was then cloned into pEGFP-N1 digested with BamHI and NotI to replace EGFP with H2B-mNG2_11_.

### Transfection

Cells were edited using transfection. Cells were treated with the NHEJ inhibitors NU7441 (2 μM; Tocris Bioscience) and SCR7 (1 μM; Xcess Biosciences) for 4 hours before and 48 hours after nucleofection, while 10 μM ROCK inhibitor was added to the media 2 hours before nucleofection to increase recovery. For AAVS1 targeting, the pAAVS1-P-CAG-mNG2_1-10_ repair template was purified prior to nucleofection using the GeneJET plasmid maxiprep kit (Thermo Fisher Scientific). Synthetic sgRNAs were purchased from MilliporeSigma and Synthego (listed in Table S3) and ArciTect Cas9-eGFP and SpCas9-2NLS were purchased from STEMCELL Technologies and Synthego, respectively. sgRNA spacer sequences were obtained from previous studies (listed in Table S3) or designed using Benchling (Doench et al., 2016; Hsu et al., 2013).

Cas9/sgRNA complexes were pre-mixed at room temperature for 15 minutes before nucleofection. ssODN repair templates were purchased from BioCorp and ThermoFisher Scientific (listed in Table S4). For AAVS1 targeting, 201B7 cells were transfected with 1 μg of repair template and 7.5 pmoles sgRNA pre-complexed with 7.5 pmoles Cas9. For endogenous tagging with mNG2_11_, cells were transfected with 82 pmoles ssODN and 91.5 pmoles sgRNA pre-complexed with 30.5 pmoles Cas9. For transient H2B-mNG2_11_ expression, cells were transfected with 750 ng of plasmid purified using the GeneJET plasmid miniprep kit (Thermo Fisher Scientific). Briefly, 5×10^5^ cells were resuspended in 20 μL P3 nucleofection buffer, mixed with the transfection reagents, and nucleofected using the DN100 program on a 4D-Nucleofector (Lonza). After nucleofection, the cells were gently resuspended in media and transferred into 6-well plates in mTeSR Plus media containing CloneR2 reagent (STEMCELL Technologies) for AAVS1 targeting, or 24-well plates containing mTeSR Plus media with 10 μM ROCK inhibitor for mNG2_11_ tagging or transient protein expression.

### Antibiotic selection

For AAVS1 integration, antibiotic selection was carried out as described in Oceguera-Yanez et al. (2016). Briefly, cells were left to recover for 3 days following transfection. On day 3, the media was changed to media with 0.5 μg/mL puromycin, which was changed every day for 10 days. Following antibiotic selection, colonies were passaged and allowed to become confluent. Edited populations were frozen and subjected to clonal isolation by FACS.

### Fluorescence-activated cell sorting, single-cell recovery and flow cytometry

FACS was used to enrich populations of edited cells or for clonal isolation. Cells were sorted using a FACSMelody cell sorter (BD Biosciences) after recovery from antibiotic selection or 8 days after nucleofection for endogenous tagging. Briefly, cells were dissociated using Accutase, resuspended in PBS (Wisent) and then passed through a 35 μm strainer to remove large cell clumps. Cells were sorted using gates set to capture individual fluorescent cells. Individual cells or enriched populations of 500 tagged cells were sorted into individual wells of a 96-well plate containing recovery media [mTeSR Plus media with CloneR2 reagent and Pen-Strep (50 units/mL Penicillin and 50 μg/mL Streptomycin; Wisent)]. To recover clones, cells were kept in recovery media for 6 days with media changes on days 2 and 4. The media was then changed daily with mTeSR Plus until day 11. On day 11, the cells were passaged into fresh 96-well plates and grown to confluency before freezing and screening.

For flow cytometry, cells were prepared and analyzed using the same protocol, and data was analyzed using the R package CytoExploreR (Hammill, 2021).

### Screening clones in 96-well plates

96-well plates containing single-cell clones were imaged on a Cytation 5 microscope (Agilent) 10 days after sorting. Briefly, 3 x 4 images were acquired for each well using a 4X phase contrast objective and stitched into one image per well. Laser autofocus was used to find the focal plane where colonies were expected to be found. Images were exported and analyzed using a custom ImageJ macro designed to identify wells with colonies. Briefly, the colony segmentation macro first crops out the well edge from the images, and identifies colonies by subtracting background signal, performing a Gaussian blur and thresholding high-contrast regions. Finally, it outlines objects and overlays them with the original well image for quality control. Finally, the macro determines object number to exclude wells with debris and/or multiple colonies and provides a list of positive and negative wells. The code for this macro is available on Github (https://github.com/CMCI/colony_screening).

### Cell lysis for PCR

Cell lysates were generated for all PCR-based assays. Edited clones grown in 96-well plates were dissociated with Accutase and split into two 96-well plates for freezing and cell lysis. The cells were washed once with PBS and resuspended thoroughly in 50 μL of QuickExtract reagent (Lucigen) per well. The resuspended cells were then transferred to PCR-compatible 96-well plates and placed at 65°C for 15 minutes followed by brief vortexing, then 98°C for 15 minutes, followed by 4°C. Lysates were stored at −20°C and thawed as needed for experiments.

### qPCR-based screening

qPCR was performed on the ViiA7 Real-Time PCR system (Applied Biosystems). The VIC-labelled RNaseP assay was purchased from ThermoFisher Scientific and used as an internal PCR control. The AmpR-specific primers and hydrolysis probe were taken from Roberts et al. (2017), and the mNG2_1-10_-specific primers and hydrolysis probe were designed as follows:

Forward primer: 5’-TACCGCTACACCTACGAGGG-3’

Reverse primer: 5’-GTCATCACAGGACCGTCAGC-3’

Probe: 5’-6-FAM/AT CAA AGG A/ZEN/G AGG CCC AGG TGA TG/IABkFQ-3’

The primers and hydrolysis probes were purchased from ThermoFisher Scientific and IDT, respectively. The AmpR and mNG2_1-10_ assays were prepared by mixing 18 μM of each primer with 5 μM of the hydrolysis probe. qPCR reactions consisted of 1 μL cell lysate, 0.5 μL RNaseP assay (ThermoFisher Scientific), 0.5 μL mNG2_1-10_ or AmpR assay, 5 μL QuantStudio 3D Digital PCR Master Mix V2 (ThermoFisher Scientific) and 3 μL water for a final volume of 10 μL. The qPCR thermocycling conditions were carried out as per manufacturer’s instructions for the QuantStudio 3D Digital PCR Master Mix V2. The data was analyzed using the ViiA7 software and clones that were positive for mNG2_1-10_ and negative for AmpR were selected for further screening by PCR as described below.

### PCR screening and sequencing

Integration of the mNG2_1-10_ cassette at the AAVS1 locus was verified by amplifying the left and right junctions of the integration, as well as the WT AAVS1 locus. Briefly, 50 μL PCR reactions were prepared as follows: 10 μL GC buffer (ThermoFisher Scientific), 1 μL 10 mM dNTP mix (ThermoFisher Scientific), 1 μL DMSO (ThermoFisher Scientific), 0.25 μL of each primer at 100 μM, 5 μL cell lysate, 0.5 μL Phusion polymerase (ThermoFisher Scientific) and 32 μL water. The primers used for each PCR reaction are listed in Table S5. The following touchdown cycles were then performed: 98°C for 3 minutes, followed by 40 cycles of: 98°C for 10 seconds, initial annealing at 72°C and decreasing by 1°C every cycle until down to 55°C, and 72°C for 30 seconds per kb; followed by a final extension at 72°C for 10 minutes and hold at 12°C. The PCR products were run on a 0.8% agarose gel stained with ethidium bromide and analyzed manually. The bands corresponding to the expected amplicons were gel extracted using the GeneJET gel extraction kit (Thermo Fisher Scientific) and sequenced by Sanger sequencing (Eurofins).

### Digital PCR

Digital PCR reactions were prepared as follow: 1.5 μL cell lysate, 8.7 μL QuantStudio 3D Digital PCR Master Mix V2 (Thermo Fisher Scientific), 0.87 μL RNaseP reference assay (Thermo Fisher Scientific), 0.87 μL mNG2_1-10_ or AmpR assay (prepared as described above), and 5.46 μL water. The amount of lysate was calculated based on the number of cells used and the dynamic range of the QuantStudio 3D system (400-4000 copies/μL), and input lysate volumes were adjusted as needed to obtain data within this dynamic range. 14.5 μL of the reaction mix was loaded onto dPCR chips using the QuantStudio 3D Digital PCR 20K Chip Kit V2 (Thermo Fisher Scientific), taken through PCR thermocycling and analyzed on the QuantStudio 3D system (Thermo Fisher Scientific) according to manufacturer’s instructions. Data was analyzed using the QuantStudio 3D AnalysisSuite Cloud Software (Thermo Fisher Scientific).

### Karyotyping

G-banding karyotyping was carried out and analyzed by the Banque de cellules leucémiques du Québec. 22 cells in metaphase were analyzed at a resolution of 400 bands per haploid karyotype and showed normal karyotypes (46, XX).

### Pluripotency marker staining

Cells were fixed and stained for common pluripotency markers to verify the absence of differentiation. smNG2-P cells were dissociated with Accutase, washed once with PBS and fixed in 4% paraformaldehyde (PFA) in PBS for 30 minutes at room temperature. To stain OCT3/4 and NANOG, the cells were washed once with PBS and treated with a permeabilization/blocking solution containing 0.1% Triton X-100 (Sigma) and 5% Normal Donkey Serum (NDS; Jackson ImmunoResearch) in PBS for 30 minutes at room temperature. To stain TRA-1-60, which is a cell surface marker, the cells were washed once and treated with a blocking solution containing 5% NDS in PBS for 30 minutes at room temperature. The following antibody dilutions were prepared in permeabilization/blocking solution or blocking solution: 1:25 anti-NANOG antibody (1 μg/mL final concentration, PCRP-NANOGP1-2D8, DSHB), 1:20 anti-OCT3/4 antibody (10 μg/mL final concentration; Santa Cruz Biotechnology), or 1:20 Alexa488-conjugated anti-TRA-1-60 antibody (7.5 μg/mL final concentration; STEMCELL Technologies). The cells were stained in 100 μL of diluted antibody per 10^6^ cells overnight at 4°C. For unconjugated primary antibodies, the cells were washed with permeabilization/blocking solution and stained with a 1:400 dilution of Alexa488-conjugated anti-mouse antibody (Invitrogen) for 1h at 4°C. The cells were then washed with permeabilization/blocking solution or blocking solution and resuspended in 1 mL PBS before flow cytometry. Cells stained with only the secondary antibody were used as a negative control.

### Trilineage differentiation and differentiation marker staining

Cells were differentiated into the three germ layers to ensure they retained pluripotent potential after editing. Directed differentiation was performed using the STEMdiff Trilineage Differentiation Kit (STEMCELL Technologies) as per manufacturer’s instructions. The following modifications were made: ectoderm differentiation was carried out on 24-well plates coated with iMatrix-511 Silk by seeding the recommended number of cells (400,000) on day 0, mesoderm differentiation was carried out on 24-well plates coated with iMatrix-511 Silk by seeding 50,000 cells per well on day 0, and endoderm differentiation was carried out on 24-well plates coated with Matrigel (Corning) by seeding 20,000 cells per well on day 0. After differentiation, the cells were harvested by dissociation with Accutase, washed with PBS, fixed with PFA and stained as described above for PAX6 (ectoderm lineage), FOXA2 (endoderm lineage) or the cell surface marker NCAM (mesoderm lineage). The following antibody dilutions were prepared in permeabilization/blocking solution or blocking solution: 1:45 anti-PAX6 antibody for ectoderm cells (1 μg/mL final concentration, PAX6, DSHB), 1:240 anti-NCAM antibody for mesoderm cells (0.25 μg/mL final concentration, 5.1H11, DSHB), 1:50 PE-conjugated anti-FOXA2 antibody (1 μg/mL final concentration; BD Biosciences) and 1:400 dilution of Alexa488-conjugated anti-mouse antibody (Invitrogen). The cells were stained in 100 μL of diluted antibody per 10^6^ cells for 1 hour at 4°C. Stained undifferentiated cells and unstained cells were used as negative controls.

### Off-target sequencing

Potential off-target sites for the AAVS1 sgRNA were selected from Wang et al. (2014), Benchling (Hsu et al., 2013) and CRISPOR (Concordet & Haeussler, 2018) based on previous studies of Cas9 or predicted off-target sites with high scores. Each off-target site was amplified as described above by touchdown Phusion PCR from 201B7 and smNG2-P cell lysates, and sequenced by Sanger sequencing (Eurofins). The primers used to amplify the 12 off-target sites were designed using Primer-BLAST (Ye et al., 2012) and are listed in Table S5. No mutations were found at the predicted off-target cut sites, and all 12 sites sequenced in the edited cell line matched the sequence from the WT cells.

### Mycoplasma test

PCR-based mycoplasma detection was performed using the Mycoplasma PCR detection kit (Applied Biological Materials) according to the manufacturer’s protocol. For each test, a positive and negative control was included. The PCR products were run on a 0.8% agarose gel stained with ethidium bromide and analyzed manually.

### Nanopore sequencing and data analysis

For Nanopore sequencing of mixed edited populations, PCR reactions were carried out as described above with a stable annealing temperature of 65°C for 30 cycles to minimize non-specific amplification and PCR bias. The samples were then gel extracted and pooled in equimolar amounts before library preparation for Nanopore sequencing.

For multiplexed clone sequencing, the edited loci from each clone were amplified individually and barcoded by PCR prior to Nanopore sequencing. For this, individual clones were subjected to a first round of PCR to amplify the target loci and add universal adapters with the primers listed in Table S5. The locus-specific primers were designed using Primer-BLAST (Ye et al., 2012) and the sequence of the universal adapters were taken from Karst et al. (2021). The first round of PCR was carried out as described above with annealing at 65°C and for 15 cycles. The PCR products were diluted 1/100 into the second PCR reactions, which contained a unique pair of barcoding primers for each clone for a specific target locus. The second PCR was carried out as described above with annealing at 65°C and for 15 cycles. The PCR products for a specific edited locus were pooled, run on a 0.8% agarose gel and gel extracted. The concentration of pooled products was then measured and the PCR products for clones edited at different target loci were then pooled in equal molar ratios before Nanopore library preparation.

Library preparation for Nanopore sequencing was carried out using the NEBNext Companion Module for Oxford Nanopore Technologies Ligation Sequencing (New England Biolabs), the Ligation Sequencing Kit V14 and the Flongle Sequencing Expansion (Oxford Nanopore Technologies), as per manufacturer’s protocols. The final library concentration was measured using a Qubit fluorometer (Invitrogen) using the Qubit dsDNA HS assay kit (Invitrogen), and 5 fmoles of the library were loaded onto a Flongle flow cell and sequenced overnight.

Basecalling of the Nanopore FAST5 files was performed with Guppy (R10.4.1 flow cell, 400 bps, super accuracy configuration) using the Compute Canada server, and low-quality reads with Q scores below 10 were excluded. Reads were demultiplexed using a custom Bowtie-based pipeline (https://github.com/frba/nanopore_demultiplex). When sequencing amplicons from mixed edited populations, reads were demultiplexed based on the presence of either of two gene-specific barcodes (listed in Table S6). The FASTQ files were analyzed using CRISPResso2 (Clement et al., 2019) to obtain the frequency of WT, HDR and indel alleles in the population. All HDR alleles were included in the same category because of the high sequencing error rate. For clone sequencing, reads were demultiplexed based on the presence of two gene-specific and two clone-specific barcodes (listed in Table S6). The FASTQ reads were aligned to the expected WT and edited sequences using Minimap2 (Li, 2018), and the alignments were visualized using IGV (Thorvaldsdottir et al., 2013). The genotypes of edited clones were inferred manually by looking at the alignments of sequencing reads with the WT and edited alleles for each clone.

### Fluorescence microscopy

The fluorescence from mNG2_1-10_/mNG2_11_ complementation was visualized the day after transfection with the pH2B-mNG2_11_ plasmid using a Cytation 5 microscope (Agilent) equipped with a 20x/0.45 NA phase contrast objective, a 465 nm LED cube and green filter cube (excitation 469/35 nm, emission 525/39 nm and dichroic mirror 497 nm).

Cells with mNG2_11_ integrated at endogenous loci were imaged at least 8 days after nucleofection on a Leica DMI6000B inverted epifluorescence microscope equipped with a EL6000 mercury lamp and a GFP3035B filter cube (excitation 472/30 nm, emission 520/35 and dichroic mirror 495 nm) using a 20x/0.35 NA objective, an Orca R2 CCD camera (Hamamatsu) and Volocity software (PerkinElmer).

Because of its weak fluorescent signal, the CDH1-mNG2_11_ edited cells were seeded a 35 mm polymer coverslip dish (Cellvis) coated with iMatrix-511 Silk, and imaged on an inverted Nikon Eclipse Ti microscope (Nikon) equipped with a Livescan Sweptfield scanner (Nikon), Piezo Z stage (Prior), IXON 879 EMCCD camera (Andor), and a 488 nm laser (50 mW, Agilent) using the 100x/1.45 NA objective.

For live imaging, cells were adapted to culture on Matrigel (Corning) for at least 2 passages. Cells were passaged as described above, and seeded onto 35 mm polymer coverslip dishes (Cellvis) freshly coated with Matrigel. The cells were then allowed to grow into colonies for 4 to 5 days before imaging. Fresh mTeSR Plus media was added to the cells at least 30 minutes before imaging. To visualize chromatin, Hoechst 33342 (Invitrogen) was added to the cells at a final concentration of 1.78 μM (1 μg/mL) for 30 minutes prior to imaging. Imaging was performed using the inverted Nikon Eclipse Ti microscope with the Livescan Sweptfield scanner described above, equipped with 405 and 488 nm lasers (50 mW, Agilent). The cells were kept at 37°C and 5% CO_2_ during imaging in an INU-TiZ-F1 chamber (MadCityLabs). Z-stacks of 9 slices at 1 μm intervals were acquired every minute using NIS Elements software (Nikon, version 4.0). For CARE training, images were collected using low- and high-exposure setting with a 16-slice Z-stack of 0.75 μm intervals. The imaging parameters (laser power and exposure time) used for each cell line in low- and high-exposure conditions are listed in Table S7.

### Image restoration by CARE

For image restoration, we used the CSBDeep neural network developed by Weigert et al. (2018). Briefly, a training dataset composed of >100 matched 2-channel fluorescent images (green for mNG and blue for Hoechst) with low- and high-exposure settings was acquired for each tagged cell line. The training datasets were designed to include cells in interphase and at different stages of mitosis. We used the Python implementation of CSBDeep to train the standard model of the neural network for each cell line individually. We then applied this model to restore timelapse Z-stack 2-channel images of the corresponding cell lines.

### Image analysis

All images acquired using NIS Elements (Nikon) were opened in Fiji (Version 2.3, NIH) for analysis. For signal-to-noise ratio measurements, two regions of interest were drawn over an area of homogeneous signal inside a cell, and over a region of background signal (in a region with no cells). The mean pixel intensity and standard deviation (SD) were measured for both regions of interest, and the signal-to-noise ratio was calculated as follows: SNR = (mean signal - mean background) / signal SD. The measurement was repeated using the same regions of interest for matched low- and high-exposure images.

Linescans were performed and measured for tagged cell lines using a macro in Fiji modified from Ozugergin et al. (2022). The macro was designed to isolate the 488 nm channel from the image file, subtract background signal and perform a bleach correction. The desired timepoint and two central Z slices were picked manually, and the macro generated a Z-stack average projection. A five pixel-wide line was then traced along the cortex of the cell, from one pole to the other, along with a straight one pixel-wide line to define the midplane. The macro then measured the fluorescence intensity of each pixel along the length of the linescan and positioned the pixels in relation to the midplane. For the measurement of tubulin at the central spindle, the same macro was used to draw a five pixel-wide line across the cell equator. All data was exported for use in Excel (Microsoft) and Prism (Version 9.3, GraphPad) for further analysis.

For breadth measurements, the number of pixels above 50% of the normalized peak intensities were counted for each linescan and converted to microns. Pixels with intensities higher than the cutoff value outside of the peak region were excluded from these calculations. For measurements of the ratio of cortical to cytosolic protein in metaphase cells, the average intensity of pixels at the cortex was measured by a linescan as described above, and the average intensity of pixels in the cytosol was measured by drawing a region of interest over the cytosol.

### Statistical analysis

Box and whiskers plots were generated using Prism (Version 9.3, GraphPad) to show median values (central line), quartiles (box edges) and minimum and maximum values (whiskers). Statistical significance was tested using a Brown-Forsythe and Welch’s ANOVA, followed by multiple comparisons using Dunnett’s T3 test, or by Welch’s t test (Graphpad Prism version 9.3). Significance levels were defined as: p>0.5 non-significant (ns), * p≤0.05; ** p≤0.01; *** p≤0.001; **** p≤0.0001.

## Acknowledgements

We thank Nicholas D. Gold and Angela Quach from the Concordia Genome Foundry for their help with equipment use and analysis of the Nanopore sequencing data. We also thank the Concordia Center for Microscopy and Cellular Imaging for help with imaging and analysis. We thank Dr. Knut Woltjen from the Center for iPS Cell Research and Application (CiRA) and his lab for guidance with iPS culture and editing, and researchers from the National Research Council of Canada (NRC) for feedback on this project. This research was funded by the NRC Disruptive Technology Solutions for Cell and Gene Therapy Challenge program and the NSERC Discovery Grant RGPIN-2023-04805. M.C.H. was supported by a FRQNT B2X and NSERC CREATE SynBioApps scholarships. A.J.P was supported by Concordia University Research Chair in Cancer Cell Biology and V.J.J.M was supported by the Applied Synthetic Biology Senior Research Chair.

---

**Figure S1.**
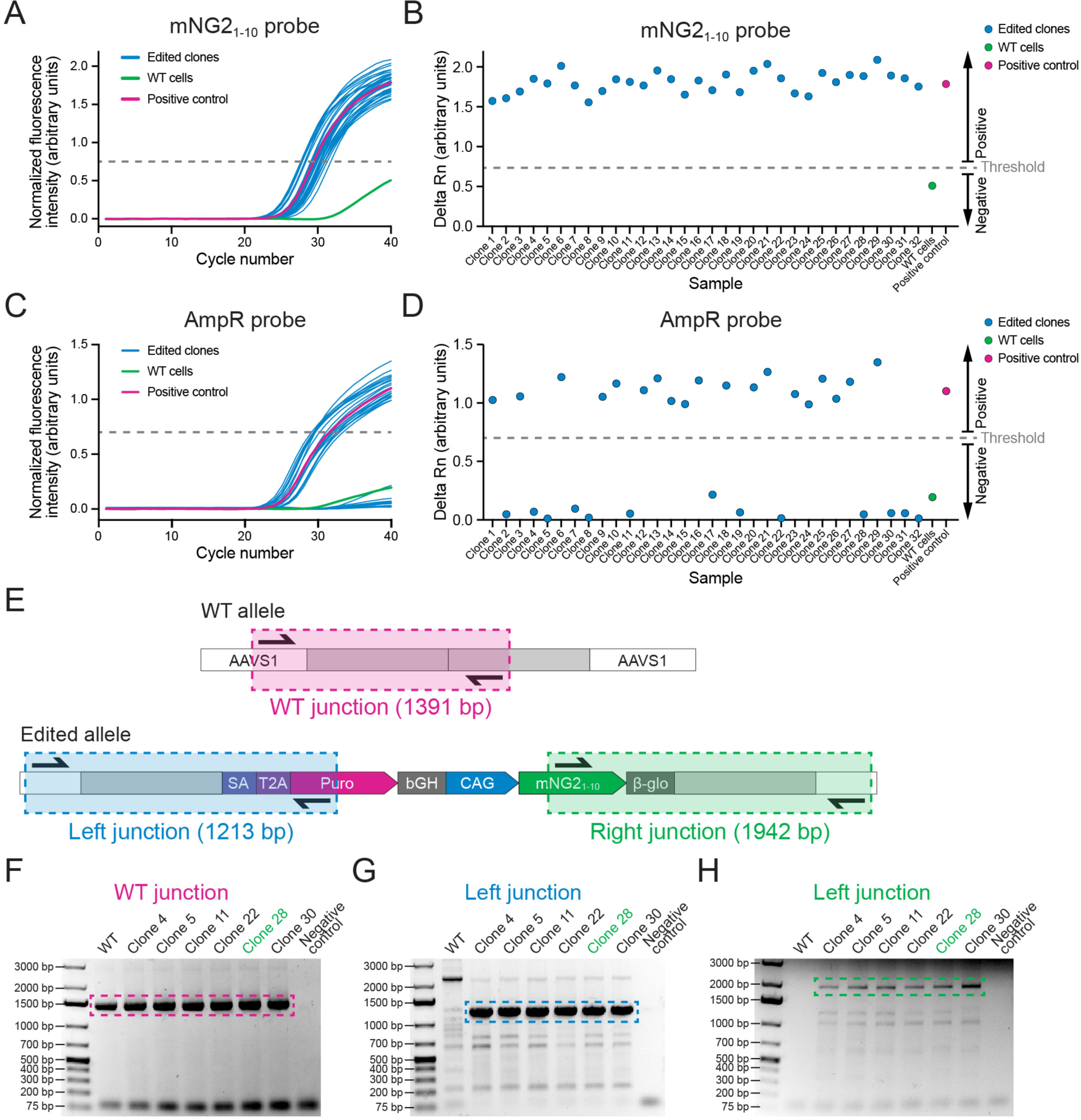
Screening of AAVS1-mNG2_1-10_ clones. **A)** A graph shows qPCR amplification curves for the mNG2_1-10_ probe across 32 edited clones (blue), WT cells (green) and a positive control (pink). **B)** A graph shows the delta Rn values (increase in fluorescence over the 40 qPCR cycles) to show the presence or absence of the mNG2_1-10_ gene in 32 clones edited clones (blue), WT cells (green) and a positive control (pink). Clones with delta Rn values above the threshold were considered positive for mNG2_1-10_. **C)** A graph shows qPCR amplification curves for the AmpR probe in 32 edited clones (blue), WT cells (green) and a positive control (pink). **D)** A graph shows the delta Rn values (increase in fluorescence over the 40 qPCR cycles) to show the presence or absence of the AmpR gene in 32 edited clones (blue), WT cells (green) and a positive control (pink). Clones with delta Rn values below threshold were considered negative for AmpR. **E)** A schematic shows the strategy for junction PCR screening. Three sets of primers were used to amplify the WT AAVS1 allele (top, pink), and the left (bottom left, blue) and right (bottom right, green) integration junctions to verify the mNG2_1-10_ insertion. **F-H)** Gel images show the results of the junction PCR screening for WT (F), left junction (G) and right junction (H) with WT cells and 6 edited clones. The hatched boxes indicate the relevant DNA band. Clone 28 (green) was selected for further validation.

**Figure S2.**
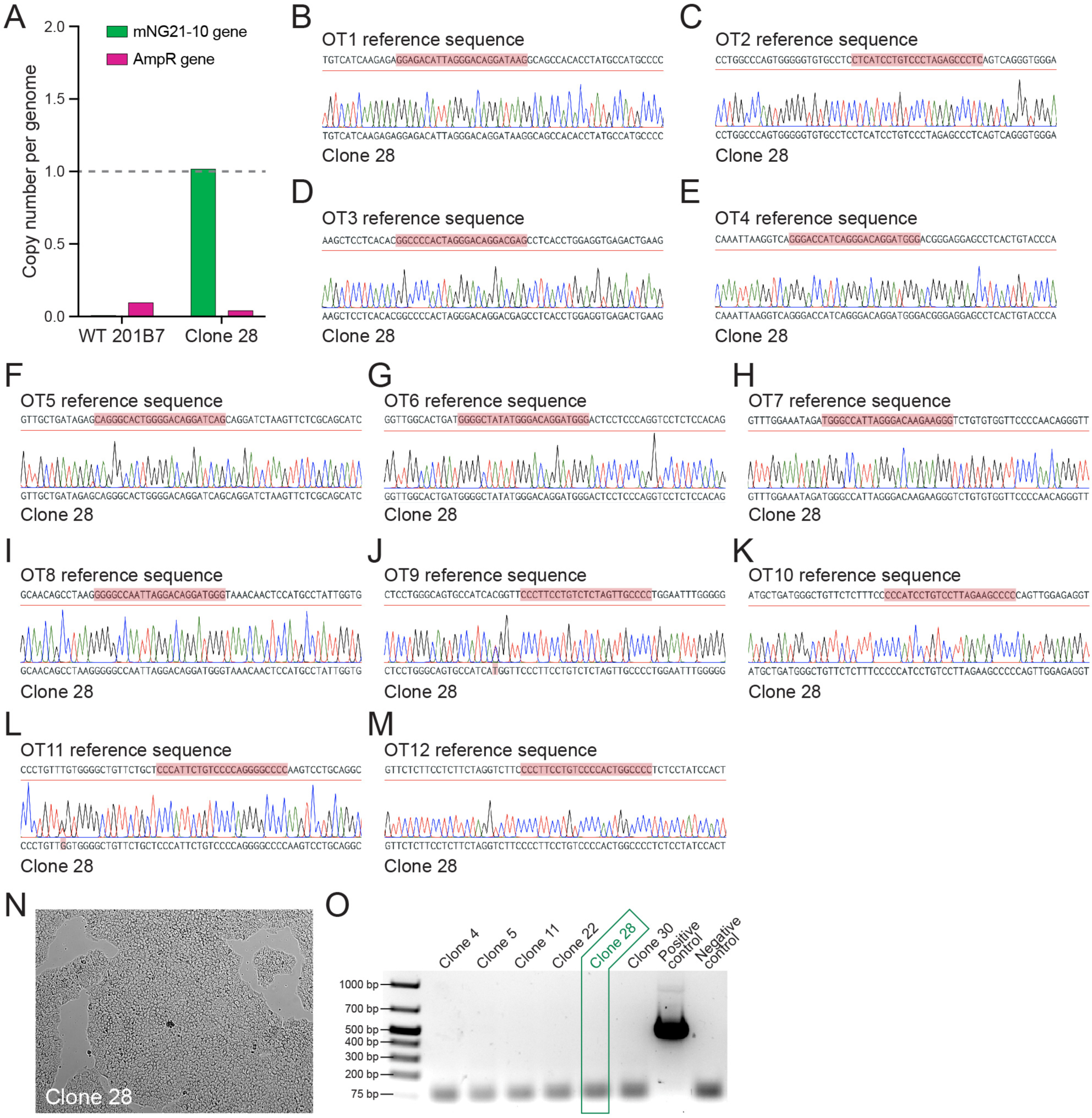
Validation of the smNG2-P cell line. **A)** A bar graph shows the copy number of the mNG2_1-10_ (green) and AmpR (pink) genes in clone 28 and WT cells determined by digital PCR. **B-M)** Sequence chromatograms for 12 AAVS1 off-target sites in AAVS1-mNG2_1-10_ clone 28 cells are shown below the reference sequence from the human genome. The mismatched sgRNA target and PAM sites are highlighted in red. **N)** A brightfield image shows AAVS1-mNG2_1-10_ clone 28 cells growing on an iMatrix-511 Silk-coated plate. **O)** A gel image shows the mycoplasma PCR test with cells from several AAVS1-mNG2_1-10_ clones, along with positive and negative controls. Clone 28 is highlighted in green.

**Figure S3.**
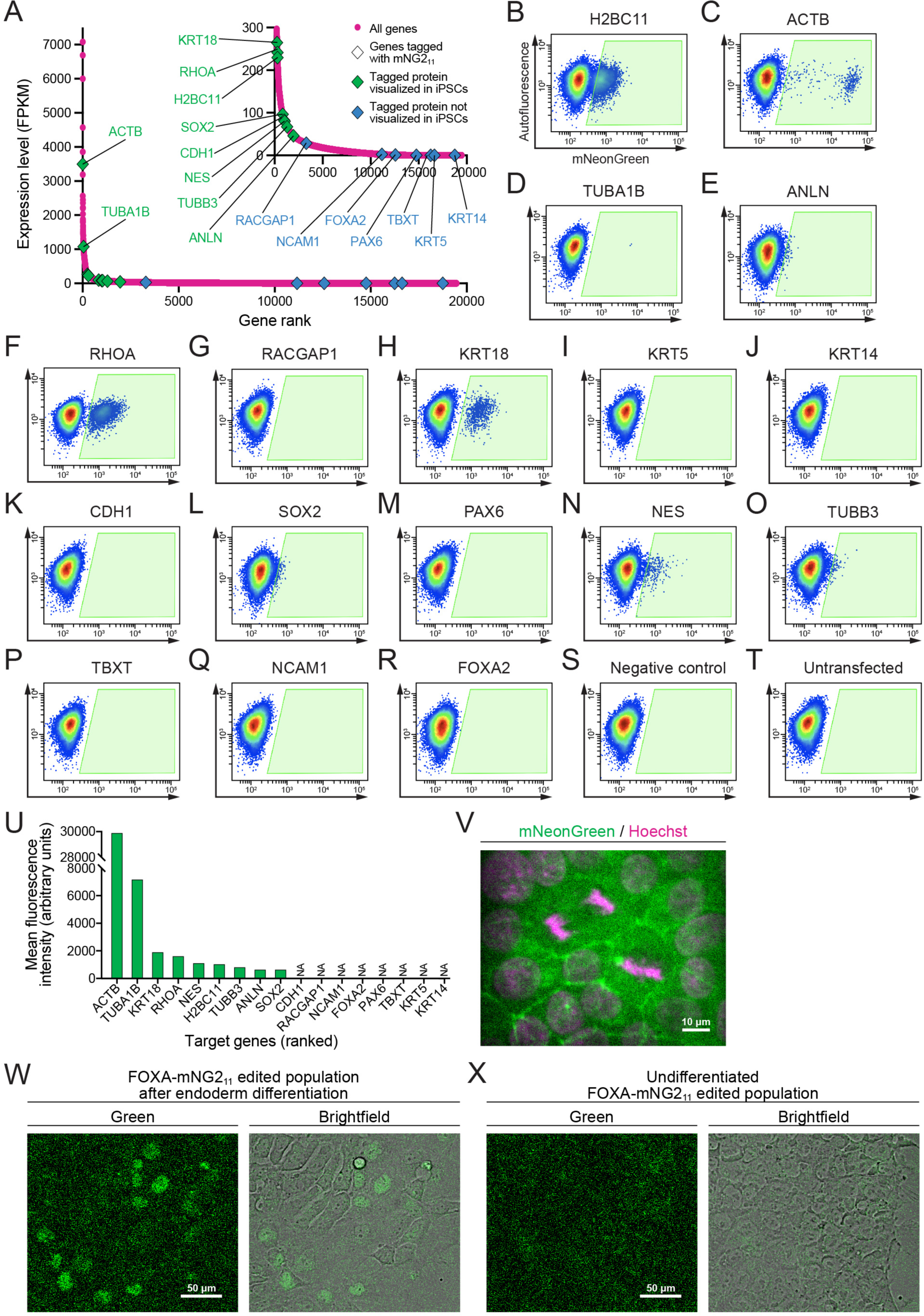
Fluorescent signal from endogenous mNG2_11_ tags. **A)** A graph shows the expression levels of protein-coding genes in the 201B7 cell line. Genes targeted for tagging with mNG2_11_ are indicated by triangles (green for tagged proteins that were detected and blue for tagged proteins that were not detected in iPSCs). The RNA-seq data was obtained from Iwasaki et al. (2022; GEO accession number GSE199820). **B-T)** Flow cytometry plots show 17 different cell populations with smNG2 fluorescence after being edited with the mNG2_11_ tag. A negative control transfected with pooled ssODNs and scrambled sgRNAs (R), and an untransfected control (S) were used to set a gate to identify fluorescent cells (gate shown in green). **U)** A bar graph shows the mean fluorescence intensity of fluorescent cells in the gates shown in (B-T). **V)** Fluorescent images show the localization of smNG2-tagged E-cadherin (green) and DNA (stained with Hoechst, magenta). The scale bar is 10 microns. **W-X)** Fluorescent and brightfield images show the expression of smNG2-tagged FOXA2 in the edited cell population after differentiation into endoderm (U) but not in the undifferentiated cell population (V). The scale bar is 50 microns.

**Figure S4.**
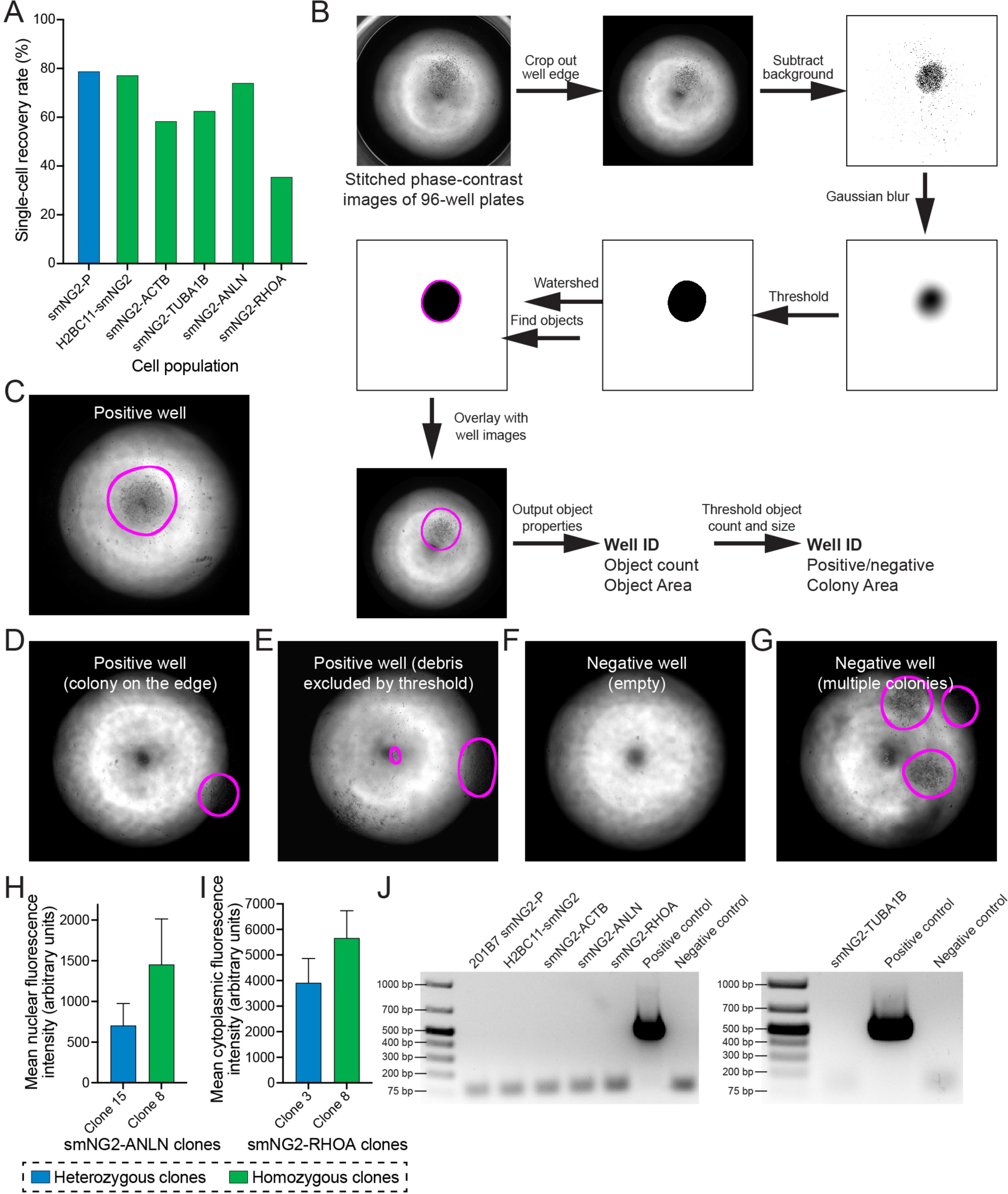
Clonal isolation and screening of tagged clones. **A)** A bar graph shows the recovery rate of different edited and WT cell populations after single-cell isolation by FACS. **B)** The steps used by the macro to process images for colony screening in 96-well plates are shown. **C-G)** Representative images of results from colony screening using the macro in B). **H)** Mean nuclear fluorescence intensity of a heterozygous (shown in blue, n=120) and homozygous (shown in green, n=112) smNG2-anillin clone measured by fluorescence microscopy. **I)** Mean cytoplasmic fluorescence intensity of a heterozygous (shown in blue, n=103) and homozygous (shown in green, n=101) smNG2-RhoA clone measured by fluorescence microscopy. **J)** Gel images show the results of the mycoplasma PCR test for the final mNG2_11_-tagged cell lines compared to positive and negative controls.

**Figure S5.**
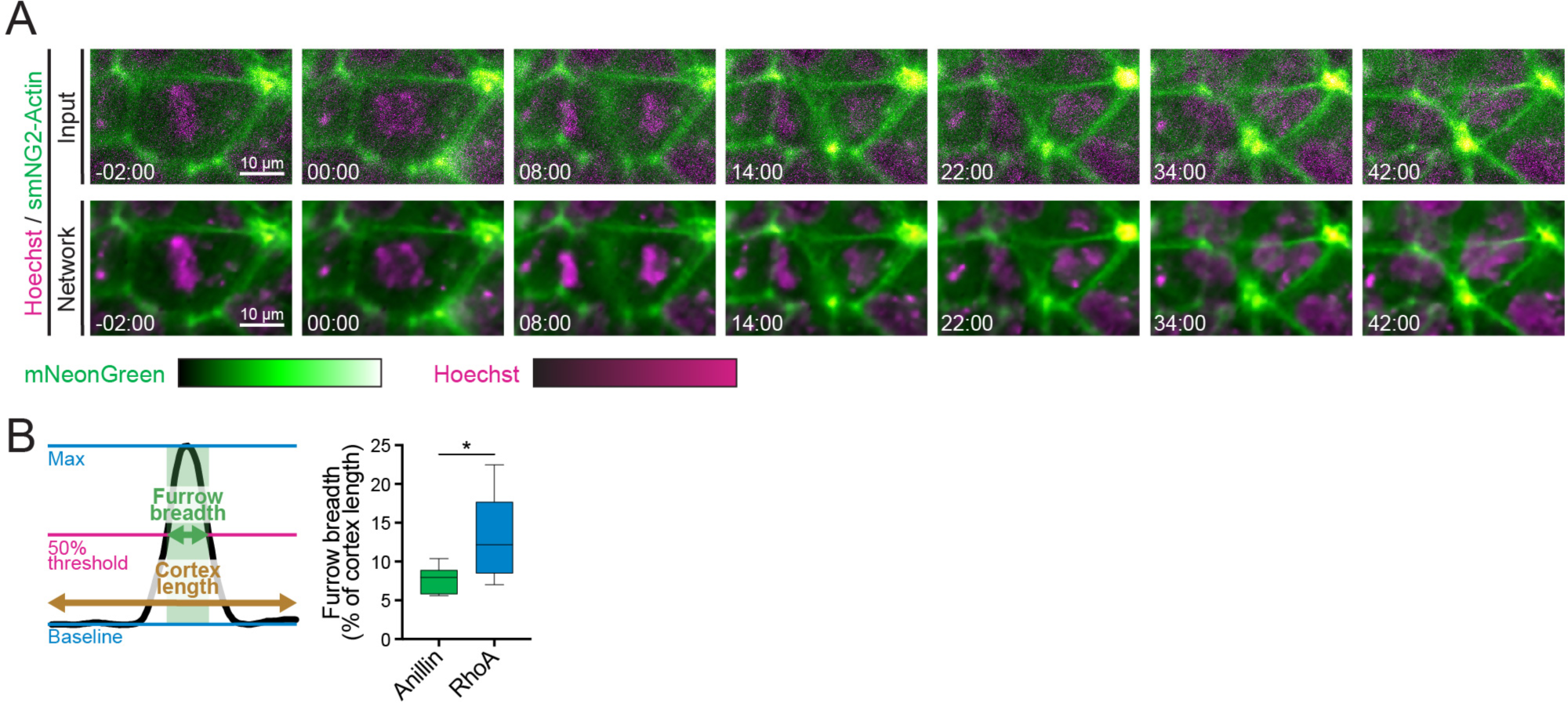
Protein localization in iPSCs during cytokinesis. **A)** Timelapse images show a cell expressing smNG2-actin during cytokinesis. The input images are shown above the same images after restoration by the CARE neural network. The mNG2 signal is shown in green and the DNA is stained with Hoechst (magenta). The scale bar is 10 microns and times are shown relative to anaphase onset (00:00). **B)** A schematic (left) shows how the breadth of protein localization at the equatorial cortex relative to cortex length was calculated for smNG2-anillin and smNG2-RhoA (right; n = 10 for each). Statistical significance was determined by Welch’s t test (ns, not significant; * p≤0.05; ** p≤0.01; *** p≤0.001; **** p≤0.0001).

**Table S1.**
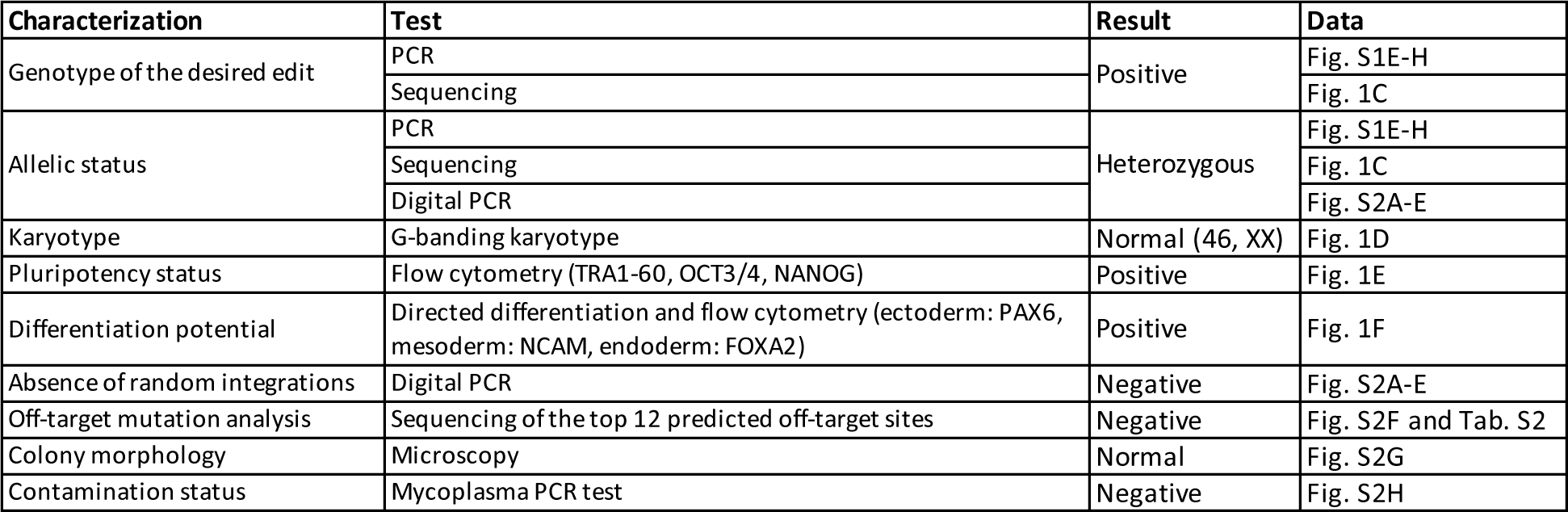
Characterization and validation of the smNG2-P cell line. The results from the validation and quality control steps carried out on the smNG2-P cell line are listed.

| Characterization | Test | Result | Data |
| --- | --- | --- | --- |
| Genotype of the desired edit | PCR | Positive | Fig. S1E-H |
|  | Sequencing |  | Fig. 1C |
| Allelic status | PCR | Heterozygous | Fig. S1E-H |
|  | Sequencing |  | Fig. 1C |
|  | Digital PCR |  | Fig. S2A-E |
| Karyotype | G-banding karyotype | Normal (46, XX) | Fig. 1D |
| Pluripotency status | Flow cytometry (TRA1-60, OCT3/4, NANOG) | Positive | Fig. 1E |
| Differentiation potential | Directed differentiation and flow cytometry (ectoderm: PAX6, mesoderm: NCAM, endoderm: FOXA2) | Positive | Fig. 1F |
| Absence of random integrations | Digital PCR | Negative | Fig. S2A-E |
| Off-target mutation analysis | Sequencing of the top 12 predicted off-target sites | Negative | Fig. S2F and Tab. S2 |
| Colony morphology | Microscopy | Normal | Fig. S2G |
| Contamination status | Mycoplasma PCR test | Negative | Fig. S2H |

**Table S2.**
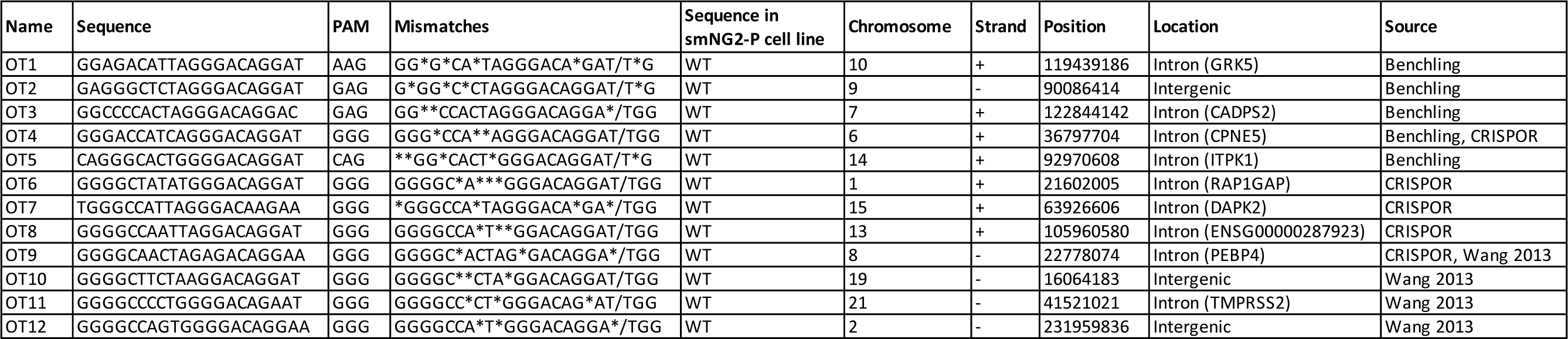
AAVS1 off-target sites sequencing in 201B7 smNG2-P cells. The top off-target sites for the AAVS1 sgRNA are listed, along with the results of sequencing these sites in the smNG2-P cell line compared to WT cells.

**Table S3.**
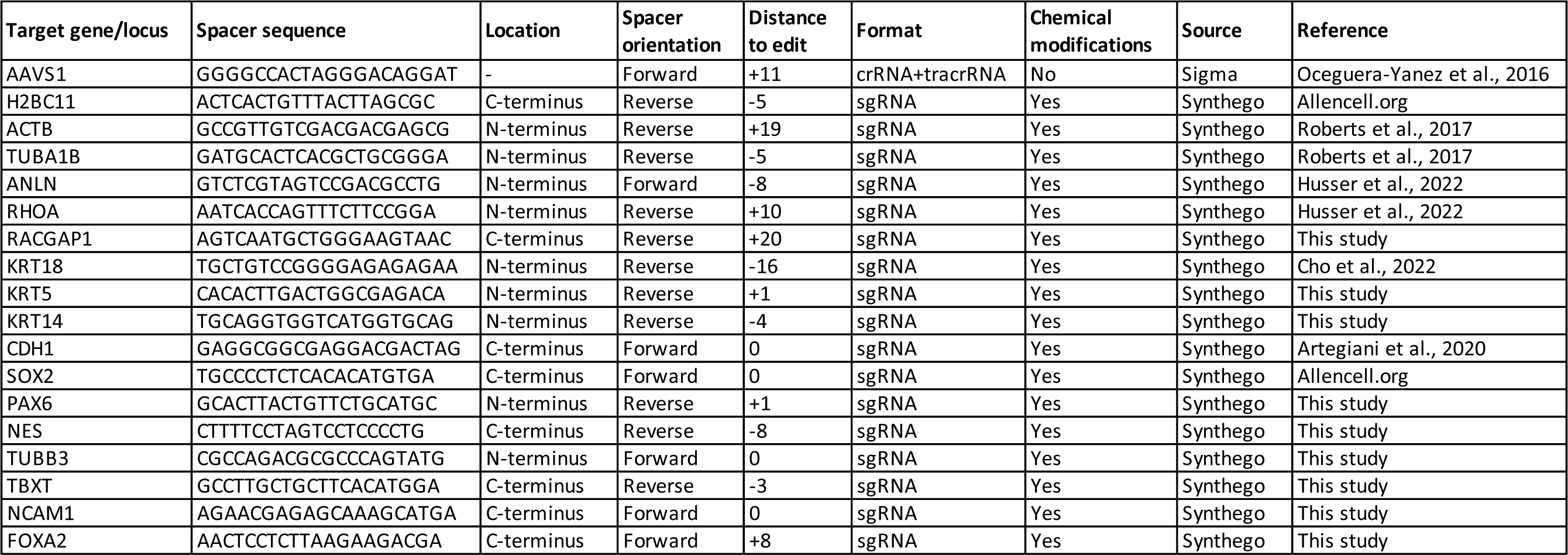
List of sgRNAs used in this study. The sequences of all sgRNAs used in this study are listed.

**Table S4.**
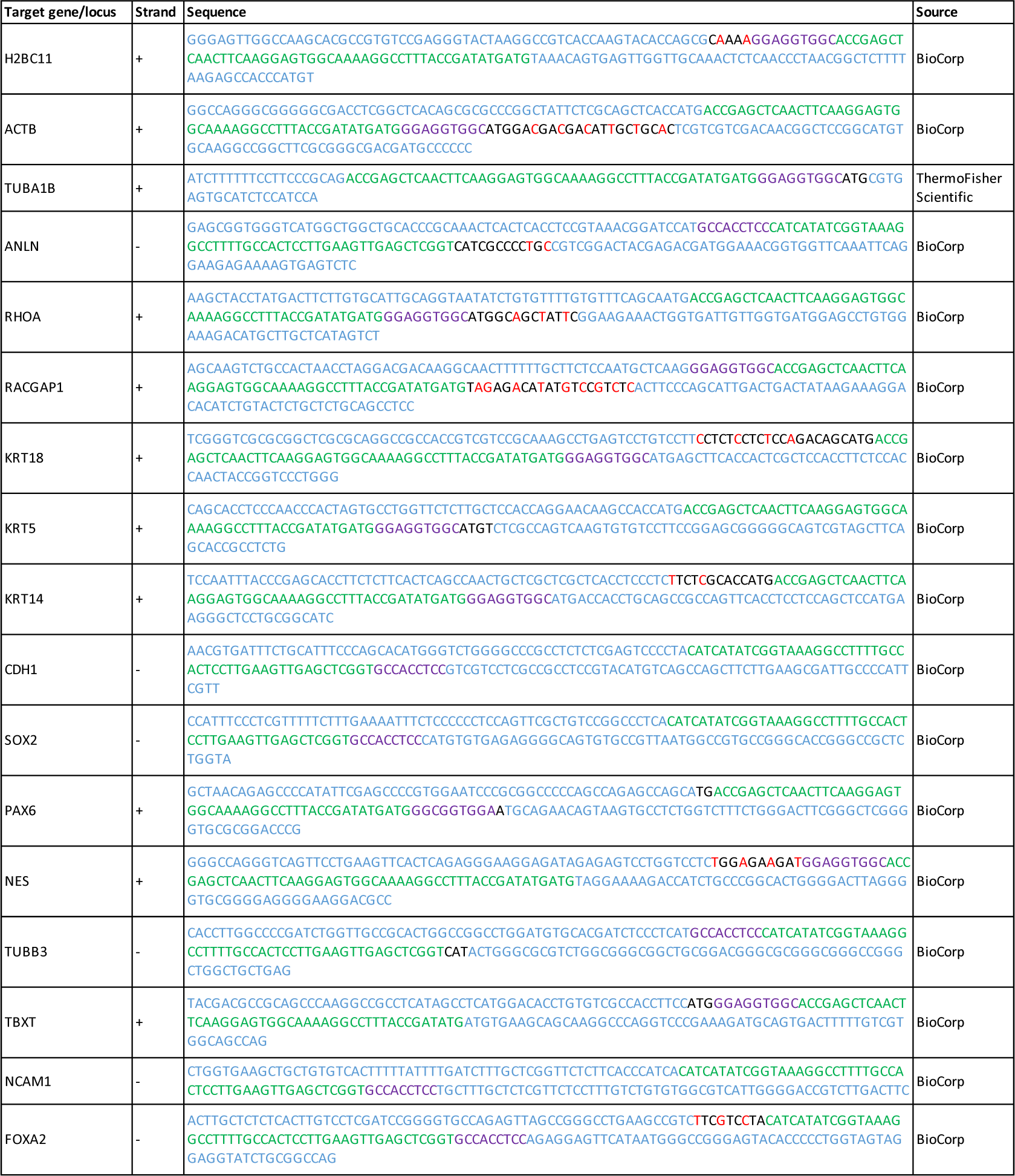
List of ssODNs used for endogenous tagging with mNG2_11_. The sequences of the single-stranded oligonucleotides used as repair templates to integrate mNG2_11_ at various loci are shown. For each oligonucleotide, the homology arms are shown in blue, the mNG2_11_ tag in green, the protein linker in purple and mutations designed to remove homology and prevent Cas9 re-cutting in red.

**Table S5.**
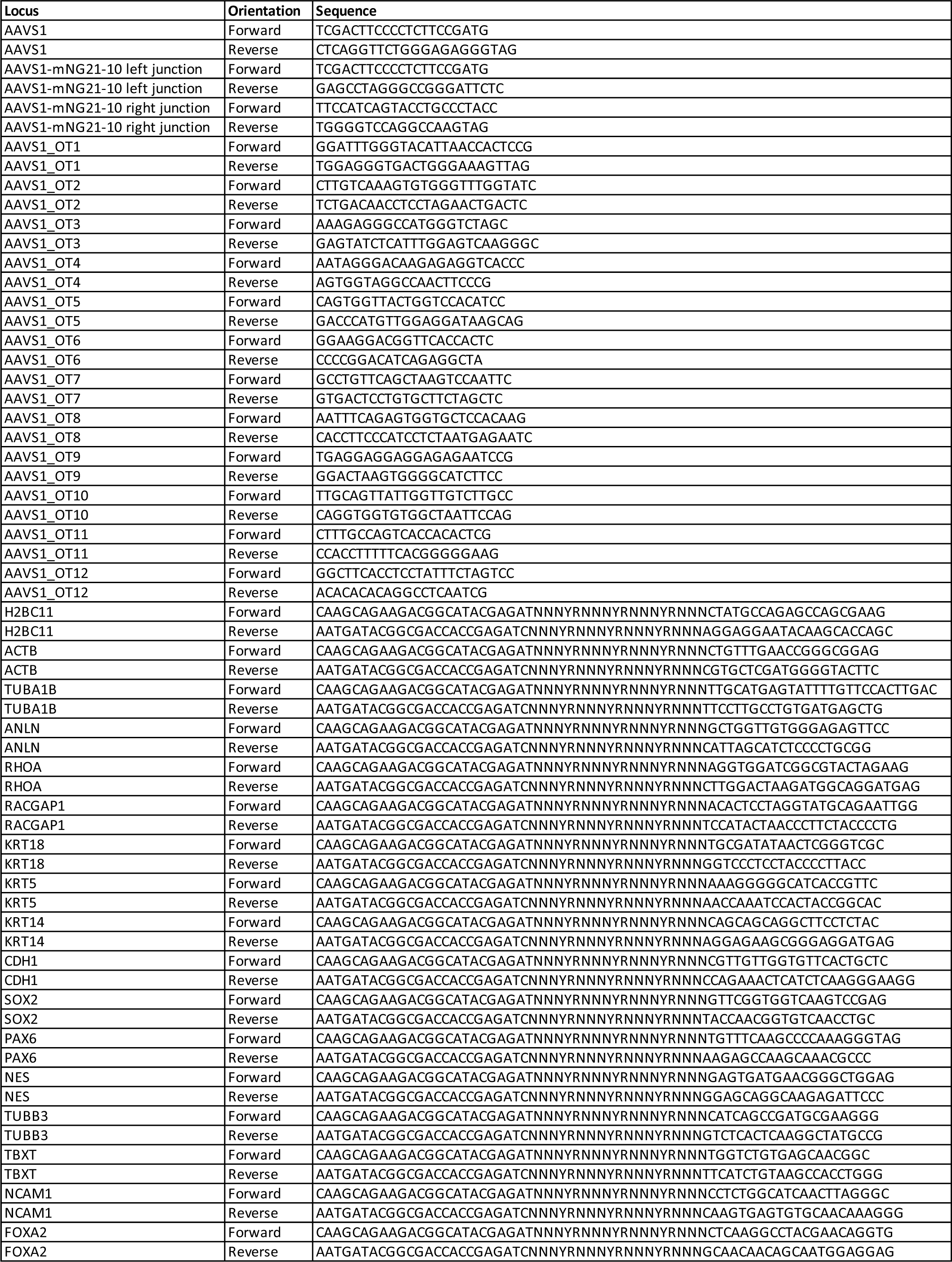
List of genotyping primers. The primers used to amplify and sequence each edited locus are listed.

**Table S6.**
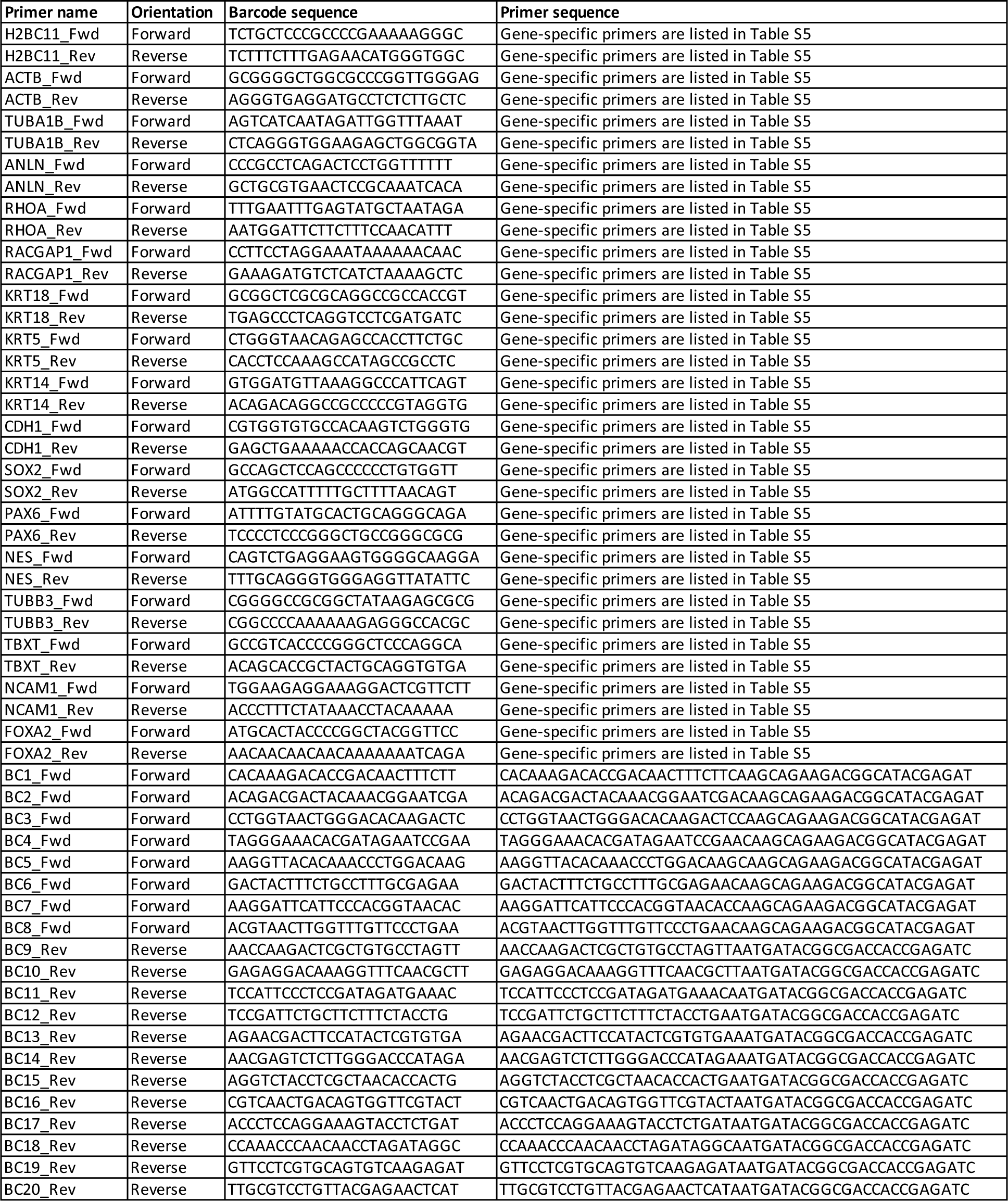
List of barcoding primers. The primers used to add unique barcodes onto locus-specific amplicons for pooled Nanopore sequencing are shown.

**Table S7.** List of live imaging parameters. For each cell lines imaged, the laser power and exposure time used to acquire low- and high-exposure images are listed. These parameters were used to acquire the neural network training dataset, and the low-exposure settings were used to acquire timelapse images for restoration.

| Cell line | Imaging condition | 405 nm laser, 50 mW |  | 488 nm laser, 50 mW |  |
| --- | --- | --- | --- | --- | --- |
|  |  | Laser power (%) | Exposure time (ms) | Laser power (%) | Exposure time (ms) |
| H2BC11-smNG2 | Low exposure | 3 | 200 | 8 | 200 |
|  | High exposure | 30 | 800 | 90 | 800 |
| smNG2-ACTB | Low exposure | 6 | 200 | 12 | 200 |
|  | High exposure | 50 | 200 | 25 | 200 |
| smNG2-TUBA1B | Low exposure | 3 | 200 | 6 | 200 |
|  | High exposure | 50 | 200 | 60 | 200 |
| smNG2-ANLN | Low exposure | 4 | 200 | 16 | 200 |
|  | High exposure | 30 | 800 | 90 | 800 |
| smNG2-RHOA | Low exposure | 4 | 200 | 20 | 200 |
|  | High exposure | 30 | 800 | 90 | 800 |

**Movie S1. *Comparison of raw (left) and restored (right) timelapse images of a smNG2-ANLN iPS cell undergoing cytokinesis.*** smNG2-anillin is shown in green and the DNA (stained with Hoechst) is shown in magenta. Images were acquired every minute and played at 4 frames per second (240x real time). Time is shown relative to anaphase onset (00:00). The scale bar is 10 microns. Still images are shown in Fig. 4C.

**Movie S2. *Comparison of raw (left) and restored (right) timelapse images of a smNG2-RHOA iPS cell undergoing cytokinesis.*** smNG2-RhoA is shown in green and the DNA (stained with Hoechst) is shown in magenta. Images were acquired every minute and played at 4 frames per second (240x real time). Time is shown relative to anaphase onset (00:00). The scale bar is 10 microns. Still images are shown in Fig. 4D.

**Movie S3. *Comparison of raw (left) and restored (right) timelapse images of a H2BC11-smNG2 iPS cell undergoing cytokinesis.*** Images were acquired every minute and played at 4 frames per second (240x real time). Time is shown relative to anaphase onset (00:00). The scale bar is 10 microns. Still images are shown in Fig. 4E.

**Movie S4. *Comparison of raw (left) and restored (right) timelapse images of a smNG2-TUBA1B iPS cell undergoing cytokinesis.*** smNG2-Tubulin is shown in green and the DNA (stained with Hoechst) is shown in magenta. Images were acquired every minute and played at 4 frames per second (240x real time). Time is shown relative to anaphase onset (00:00). The scale bar is 10 microns. Still images are shown in Fig. 4F.

**Movie S5. *Comparison of raw (left) and restored (right) timelapse images of a smNG2-ACTB iPS cell undergoing cytokinesis.*** smNG2-Actin is shown in green and the DNA (stained with Hoechst) is shown in magenta. Images were acquired every minute and played at 4 frames per second (240x real time). Time is shown relative to anaphase onset (00:00). The scale bar is 10 microns. Still images are shown in Fig. 4G.

**Movie S6. *Comparison of raw (left) and restored (right) timelapse images of a smNG2-ACTB iPS cell showing colony-scale re-arrangements of the actin network after cytokinesis.*** smNG2-Actin is shown in green and the DNA (stained with Hoechst) is shown in magenta. Images were acquired every minute and played at 4 frames per second (240x real time). Time is shown relative to anaphase onset (00:00). The scale bar is 10 microns. Still images are shown in Fig. S5A.

## Notes

### Competing Interest Statement

The authors have declared no competing interest.

### Summary of Updates

Added Figures S3U and S4H-I; updated the results section; added supplementary movies.

## References

Aban, C. E., Lombardi, A., Neiman, G., Biani, M. C., La Greca, A., Waisman, A., Moro, L. N., Sevlever, G., Miriuka, S., & Luzzani, C. (2021, Jan 21). Downregulation of E-cadherin in pluripotent stem cells triggers partial EMT. Sci Rep, 11(1), 2048. 10.1038/s41598-021-81735-1

Adriaans, I. E., Basant, A., Ponsioen, B., Glotzer, M., & Lens, S. M. A. (2019, Apr 1). PLK1 plays dual roles in centralspindlin regulation during cytokinesis. J Cell Biol, 218(4), 1250–1264. 10.1083/jcb.201805036

Beaudet, D., Akhshi, T., Phillipp, J., Law, C., & Piekny, A. (2017, Nov 15). Active Ran regulates anillin function during cytokinesis. Mol Biol Cell, 28(24), 3517–3531. 10.1091/mbc.E17-04-0253

Bhargava, N., Thakur, P., Muruganandam, T. P., Jaitly, S., Gupta, P., Lohani, N., Goswami, S. G., Saravanakumar, V., Bhattacharya, S. K., Jain, S., & Ramalingam, S. (2022, Aug). Development of an efficient single-cell cloning and expansion strategy for genome edited induced pluripotent stem cells. Mol Biol Rep, 49(8), 7887–7898. 10.1007/s11033-022-07621-9

Bukhari, H., & Muller, T. (2019, Nov). Endogenous Fluorescence Tagging by CRISPR. Trends Cell Biol, 29(11), 912–928. 10.1016/j.tcb.2019.08.004

Burgio, G., & Teboul, L. (2020, Dec). Anticipating and Identifying Collateral Damage in Genome Editing. Trends Genet, 36(12), 905–914. 10.1016/j.tig.2020.09.011

Cabrera, A., Edelstein, H. I., Glykofrydis, F., Love, K. S., Palacios, S., Tycko, J., Zhang, M., Lensch, S., Shields, C. E., Livingston, M., Weiss, R., Zhao, H., Haynes, K. A., Morsut, L., Chen, Y. Y., Khalil, A. S., Wong, W. W., Collins, J. J., Rosser, S. J., Polizzi, K., Elowitz, M. B., Fussenegger, M., Hilton, I. B., Leonard, J. N., Bintu, L., Galloway, K. E., & Deans, T. L. (2022, Dec 21). The sound of silence: Transgene silencing in mammalian cell engineering. Cell Syst, 13(12), 950–973. 10.1016/j.cels.2022.11.005

Chaigne, A., Labouesse, C., White, I. J., Agnew, M., Hannezo, E., Chalut, K. J., & Paluch, E. K. (2020, Oct 26). Abscission Couples Cell Division to Embryonic Stem Cell Fate. Dev Cell, 55(2), 195–208 e195. 10.1016/j.devcel.2020.09.001

Chaigne, A., Smith, M. B., Lopez Cavestany, R., Hannezo, E., Chalut, K. J., & Paluch, E. K. (2021, Jul 15). Three-dimensional geometry controls division symmetry in stem cell colonies. J Cell Sci, 134(14). 10.1242/jcs.255018

Chalfie, M., Tu, Y., Euskirchen, G., Ward, W. W., & Prasher, D. C. (1994, Feb 11). Green fluorescent protein as a marker for gene expression. Science, 263(5148), 802–805. 10.1126/science.8303295

Chen, Y. H., & Pruett-Miller, S. M. (2018, Aug). Improving single-cell cloning workflow for gene editing in human pluripotent stem cells. Stem Cell Res, 31, 186–192. 10.1016/j.scr.2018.08.003

Cho, N. H., Cheveralls, K. C., Brunner, A. D., Kim, K., Michaelis, A. C., Raghavan, P., Kobayashi, H., Savy, L., Li, J. Y., Canaj, H., Kim, J. Y. S., Stewart, E. M., Gnann, C., McCarthy, F., Cabrera, J. P., Brunetti, R. M., Chhun, B. B., Dingle, G., Hein, M. Y., Huang, B., Mehta, S. B., Weissman, J. S., Gomez-Sjoberg, R., Itzhak, D. N., Royer, L. A., Mann, M., & Leonetti, M. D. (2022, Mar 11). OpenCell: Endogenous tagging for the cartography of human cellular organization. Science, 375(6585), eabi6983. 10.1126/science.abi6983

Chu, V. T., Weber, T., Wefers, B., Wurst, W., Sander, S., Rajewsky, K., & Kuhn, R. (2015, May). Increasing the efficiency of homology-directed repair for CRISPR-Cas9-induced precise gene editing in mammalian cells. Nat Biotechnol, 33(5), 543–548. 10.1038/nbt.3198

Clavel, D., Gotthard, G., von Stetten, D., De Sanctis, D., Pasquier, H., Lambert, G. G., Shaner, N. C., & Royant, A. (2016, Dec 1). Structural analysis of the bright monomeric yellow-green fluorescent protein mNeonGreen obtained by directed evolution. Acta Crystallogr D Struct Biol, 72(Pt 12), 1298–1307. 10.1107/S2059798316018623

Clement, K., Rees, H., Canver, M. C., Gehrke, J. M., Farouni, R., Hsu, J. Y., Cole, M. A., Liu, D. R., Joung, J. K., Bauer, D. E., & Pinello, L. (2019, Mar). CRISPResso2 provides accurate and rapid genome editing sequence analysis. Nat Biotechnol, 37(3), 224–226. 10.1038/s41587-019-0032-3

Concordet, J. P., & Haeussler, M. (2018, Jul 2). CRISPOR: intuitive guide selection for CRISPR/Cas9 genome editing experiments and screens. Nucleic Acids Res, 46(W1), W242–W245. 10.1093/nar/gky354

Cumin, C., Huang, Y. L., Rossdam, C., Ruoff, F., Cespedes, S. P., Liang, C. Y., Lombardo, F. C., Coelho, R., Rimmer, N., Konantz, M., Lopez, M. N., Alam, S., Schmidt, A., Calabrese, D., Fedier, A., Vlajnic, T., von Itzstein, M., Templin, M., Buettner, F. F. R., Everest-Dass, A., Heinzelmann-Schwarz, V., & Jacob, F. (2022, Aug 16). Glycosphingolipids are mediators of cancer plasticity through independent signaling pathways. Cell Rep, 40(7), 111181. 10.1016/j.celrep.2022.111181

Dambournet, D., Sochacki, K. A., Cheng, A. T., Akamatsu, M., Taraska, J. W., Hockemeyer, D., & Drubin, D. G. (2018, Sep 3). Genome-edited human stem cells expressing fluorescently labeled endocytic markers allow quantitative analysis of clathrin-mediated endocytosis during differentiation. J Cell Biol, 217(9), 3301–3311. 10.1083/jcb.201710084

Doench, J. G., Fusi, N., Sullender, M., Hegde, M., Vaimberg, E. W., Donovan, K. F., Smith, I., Tothova, Z., Wilen, C., Orchard, R., Virgin, H. W., Listgarten, J., & Root, D. E. (2016, Feb). Optimized sgRNA design to maximize activity and minimize off-target effects of CRISPR-Cas9. Nat Biotechnol, 34(2), 184–191. 10.1038/nbt.3437

Doyon, J. B., Zeitler, B., Cheng, J., Cheng, A. T., Cherone, J. M., Santiago, Y., Lee, A. H., Vo, T. D., Doyon, Y., Miller, J. C., Paschon, D. E., Zhang, L., Rebar, E. J., Gregory, P. D., Urnov, F. D., & Drubin, D. G. (2011, Mar). Rapid and efficient clathrin-mediated endocytosis revealed in genome-edited mammalian cells. Nat Cell Biol, 13(3), 331–337. 10.1038/ncb2175

Drubin, D. G., & Hyman, A. A. (2017, Jun 1). Stem cells: the new “model organism”. Mol Biol Cell, 28(11), 1409–1411. 10.1091/mbc.E17-03-0183

Feng, S., Sekine, S., Pessino, V., Li, H., Leonetti, M. D., & Huang, B. (2017, Aug 29). Improved split fluorescent proteins for endogenous protein labeling. Nat Commun, 8(1), 370. 10.1038/s41467-017-00494-8

Feng, S., Varshney, A., Coto Villa, D., Modavi, C., Kohler, J., Farah, F., Zhou, S., Ali, N., Muller, J. D., Van Hoven, M. K., & Huang, B. (2019). Bright split red fluorescent proteins for the visualization of endogenous proteins and synapses. Commun Biol, 2, 344. 10.1038/s42003-019-0589-x

Gibson, T. J., Seiler, M., & Veitia, R. A. (2013, Aug). The transience of transient overexpression. Nat Methods, 10(8), 715–721. 10.1038/nmeth.2534

Grancharova, T., Gerbin, K. A., Rosenberg, A. B., Roco, C. M., Arakaki, J. E., DeLizo, C. M., Dinh, S. Q., Donovan-Maiye, R. M., Hirano, M., Nelson, A. M., Tang, J., Theriot, J. A., Yan, C., Menon, V., Palecek, S. P., Seelig, G., & Gunawardane, R. N. (2021, Aug 4). A comprehensive analysis of gene expression changes in a high replicate and open-source dataset of differentiating hiPSC-derived cardiomyocytes. Sci Rep, 11(1), 15845. 10.1038/s41598-021-94732-1

Green, R. A., Paluch, E., & Oegema, K. (2012). Cytokinesis in Animal Cells. Annual Review of Cell and Developmental Biology, Vol 28, 28, 29-+. 10.1146/annurev-cellbio-101011-155718

Guo, J., Walss-Bass, C., & Luduena, R. F. (2010, Jul). The beta isotypes of tubulin in neuronal differentiation. Cytoskeleton (Hoboken), 67(7), 431–441. 10.1002/cm.20455

Hammill, D. (2021). CytoExploreR: Interactive Analysis of Cytometry Data. R package version 1.1.0. Github. https://github.com/DillonHammill/CytoExploreR

Hesse, M., Raulf, A., Pilz, G. A., Haberlandt, C., Klein, A. M., Jabs, R., Zaehres, H., Fugemann, C. J., Zimmermann, K., Trebicka, J., Welz, A., Pfeifer, A., Roll, W., Kotlikoff, M. I., Steinhauser, C., Gotz, M., Scholer, H. R., & Fleischmann, B. K. (2012). Direct visualization of cell division using high-resolution imaging of M-phase of the cell cycle. Nat Commun, 3, 1076. 10.1038/ncomms2089

Hong, Y. J., & Do, J. T. (2019). Neural Lineage Differentiation From Pluripotent Stem Cells to Mimic Human Brain Tissues. Front Bioeng Biotechnol, 7, 400. 10.3389/fbioe.2019.00400

Hsu, P. D., Scott, D. A., Weinstein, J. A., Ran, F. A., Konermann, S., Agarwala, V., Li, Y., Fine, E. J., Wu, X., Shalem, O., Cradick, T. J., Marraffini, L. A., Bao, G., & Zhang, F. (2013, Sep). DNA targeting specificity of RNA-guided Cas9 nucleases. Nat Biotechnol, 31(9), 827–832. 10.1038/nbt.2647

Huh, W. K., Falvo, J. V., Gerke, L. C., Carroll, A. S., Howson, R. W., Weissman, J. S., & O’Shea, E. K. (2003, Oct 16). Global analysis of protein localization in budding yeast. Nature, 425(6959), 686–691. 10.1038/nature02026

Husser, M. C., Ozugergin, I., Resta, T., Martin, V. J. J., & Piekny, A. J. (2022, Nov). Cytokinetic diversity in mammalian cells is revealed by the characterization of endogenous anillin, Ect2 and RhoA. Open Biol, 12(11), 220247. 10.1098/rsob.220247

Husser, M. C., Skaik, N., Martin, V. J. J., & Piekny, A. (2021, Apr 15). CRISPR-Cas tools to study gene function in cytokinesis. J Cell Sci, 134(8). 10.1242/jcs.254409

Iwasaki, M., Kawahara, Y., Okubo, C., Yamakawa, T., Nakamura, M., Tabata, T., Nishi, Y., Narita, M., Ohta, A., Saito, H., Yamamoto, T., Nakagawa, M., Yamanaka, S., & Takahashi, K. (2022, May 20). Multi-omics approach reveals posttranscriptionally regulated genes are essential for human pluripotent stem cells. iScience, 25(5), 104289. 10.1016/j.isci.2022.104289

Kamiyama, D., Sekine, S., Barsi-Rhyne, B., Hu, J., Chen, B., Gilbert, L. A., Ishikawa, H., Leonetti, M. D., Marshall, W. F., Weissman, J. S., & Huang, B. (2016, Mar 18). Versatile protein tagging in cells with split fluorescent protein. Nat Commun, 7, 11046. 10.1038/ncomms11046

Karst, S. M., Ziels, R. M., Kirkegaard, R. H., Sorensen, E. A., McDonald, D., Zhu, Q., Knight, R., & Albertsen, M. (2021, Feb). High-accuracy long-read amplicon sequences using unique molecular identifiers with Nanopore or PacBio sequencing. Nat Methods, 18(2), 165–169. 10.1038/s41592-020-01041-y

Khaliullin, R. N., Green, R. A., Shi, L. Z., Gomez-Cavazos, J. S., Berns, M. W., Desai, A., & Oegema, K. (2018, Jul 2). A positive-feedback-based mechanism for constriction rate acceleration during cytokinesis in Caenorhabditis elegans. Elife, 7. 10.7554/eLife.36073

Kim, S., Kim, D., Cho, S. W., Kim, J., & Kim, J. S. (2014, Jun). Highly efficient RNA-guided genome editing in human cells via delivery of purified Cas9 ribonucleoproteins. Genome Res, 24(6), 1012–1019. 10.1101/gr.171322.113

Koh, S. P., Pham, N. P., & Piekny, A. (2022, Jan). Seeing is believing: tools to study the role of Rho GTPases during cytokinesis. Small GTPases, 13(1), 211–224. 10.1080/21541248.2021.1957384

Koker, T., Fernandez, A., & Pinaud, F. (2018, Mar 28). Characterization of Split Fluorescent Protein Variants and Quantitative Analyses of Their Self-Assembly Process. Sci Rep, 8(1), 5344. 10.1038/s41598-018-23625-7

Kotynkova, K., Su, K. C., West, S. C., & Petronczki, M. (2016, Dec 6). Plasma Membrane Association but Not Midzone Recruitment of RhoGEF ECT2 Is Essential for Cytokinesis. Cell Rep, 17(10), 2672–2686. 10.1016/j.celrep.2016.11.029

Kuang, Y. L., Munoz, A., Nalula, G., Santostefano, K. E., Sanghez, V., Sanchez, G., Terada, N., Mattis, A. N., Iacovino, M., Iribarren, C., Krauss, R. M., & Medina, M. W. (2019, May). Evaluation of commonly used ectoderm markers in iPSC trilineage differentiation. Stem Cell Res, 37, 101434. 10.1016/j.scr.2019.101434

Lee, M. E., DeLoache, W. C., Cervantes, B., & Dueber, J. E. (2015, Sep 18). A Highly Characterized Yeast Toolkit for Modular, Multipart Assembly. ACS Synth Biol, 4(9), 975–986. 10.1021/sb500366v

Leite, J., Chan, F. Y., Osorio, D. S., Saramago, J., Sobral, A. F., Silva, A. M., Gassmann, R., & Carvalho, A. X. (2020). Equatorial Non-muscle Myosin II and Plastin Cooperate to Align and Compact F-actin Bundles in the Cytokinetic Ring. Front Cell Dev Biol, 8, 573393. 10.3389/fcell.2020.573393

Leonetti, M. D., Sekine, S., Kamiyama, D., Weissman, J. S., & Huang, B. (2016, Jun 21). A scalable strategy for high-throughput GFP tagging of endogenous human proteins. Proc Natl Acad Sci U S A, 113(25), E3501–3508. 10.1073/pnas.1606731113

Li, H. (2018, Sep 15). Minimap2: pairwise alignment for nucleotide sequences. Bioinformatics, 34(18), 3094–3100. 10.1093/bioinformatics/bty191

Liang, X., Potter, J., Kumar, S., Zou, Y., Quintanilla, R., Sridharan, M., Carte, J., Chen, W., Roark, N., Ranganathan, S., Ravinder, N., & Chesnut, J. D. (2015, Aug 20). Rapid and highly efficient mammalian cell engineering via Cas9 protein transfection. J Biotechnol, 208, 44–53. 10.1016/j.jbiotec.2015.04.024

Luo, Y., Liu, C., Cerbini, T., San, H., Lin, Y., Chen, G., Rao, M. S., & Zou, J. (2014, Jul). Stable enhanced green fluorescent protein expression after differentiation and transplantation of reporter human induced pluripotent stem cells generated by AAVS1 transcription activator-like effector nucleases. Stem Cells Transl Med, 3(7), 821–835. 10.5966/sctm.2013-0212

Mahdessian, D., Cesnik, A. J., Gnann, C., Danielsson, F., Stenstrom, L., Arif, M., Zhang, C., Le, T., Johansson, F., Schutten, R., Backstrom, A., Axelsson, U., Thul, P., Cho, N. H., Carja, O., Uhlen, M., Mardinoglu, A., Stadler, C., Lindskog, C., Ayoglu, B., Leonetti, M. D., Ponten, F., Sullivan, D. P., & Lundberg, E. (2021, Feb). Spatiotemporal dissection of the cell cycle with single-cell proteogenomics. Nature, 590(7847), 649–654. 10.1038/s41586-021-03232-9

Mahen, R., Koch, B., Wachsmuth, M., Politi, A. Z., Perez-Gonzalez, A., Mergenthaler, J., Cai, Y., & Ellenberg, J. (2014, Nov 5). Comparative assessment of fluorescent transgene methods for quantitative imaging in human cells. Mol Biol Cell, 25(22), 3610–3618. 10.1091/mbc.E14-06-1091

Mahlandt, E. K., Arts, J. J. G., van der Meer, W. J., van der Linden, F. H., Tol, S., van Buul, J. D., Gadella, T. W. J., & Goedhart, J. (2021, Sep 1). Visualizing endogenous Rho activity with an improved localization-based, genetically encoded biosensor. J Cell Sci, 134(17). 10.1242/jcs.258823

Maruyama, T., Dougan, S. K., Truttmann, M. C., Bilate, A. M., Ingram, J. R., & Ploegh, H. L. (2015, May). Increasing the efficiency of precise genome editing with CRISPR-Cas9 by inhibition of nonhomologous end joining. Nat Biotechnol, 33(5), 538–542. 10.1038/nbt.3190

Maurer, J., Nelson, B., Cecena, G., Bajpai, R., Mercola, M., Terskikh, A., & Oshima, R. G. (2008). Contrasting expression of keratins in mouse and human embryonic stem cells. PLoS One, 3(10), e3451. 10.1371/journal.pone.0003451

Maurissen, T. L., & Woltjen, K. (2020, Jun 8). Synergistic gene editing in human iPS cells via cell cycle and DNA repair modulation. Nat Commun, 11(1), 2876. 10.1038/s41467-020-16643-5

O’Hagan, D., Kruger, R. E., Gu, B., & Ralston, A. (2021, Jul 1). Efficient generation of endogenous protein reporters for mouse development. Development, 148(13). 10.1242/dev.197418

Oceguera-Yanez, F., Avila-Robinson, A., & Woltjen, K. (2022). Differentiation of pluripotent stem cells for modeling human skin development and potential applications. Front Cell Dev Biol, 10, 1030339. 10.3389/fcell.2022.1030339

Oceguera-Yanez, F., Kim, S. I., Matsumoto, T., Tan, G. W., Xiang, L., Hatani, T., Kondo, T., Ikeya, M., Yoshida, Y., Inoue, H., & Woltjen, K. (2016, May 15). Engineering the AAVS1 locus for consistent and scalable transgene expression in human iPSCs and their differentiated derivatives. Methods, 101, 43–55. 10.1016/j.ymeth.2015.12.012

Osorio, D. S., Chan, F. Y., Saramago, J., Leite, J., Silva, A. M., Sobral, A. F., Gassmann, R., & Carvalho, A. X. (2019, Nov 12). Crosslinking activity of non-muscle myosin II is not sufficient for embryonic cytokinesis in C. elegans. Development, 146(21). 10.1242/dev.179150

Ozugergin, I., Mastronardi, K., Law, C., & Piekny, A. (2022, Jan 13). Diverse mechanisms regulate contractile ring assembly for cytokinesis in the two-cell C. elegans embryo. J Cell Sci. 10.1242/jcs.258921

Ozugergin, I., & Piekny, A. (2022). Diversity is the spice of life: An overview of how cytokinesis regulation varies with cell type. Front Cell Dev Biol, 10, 1007614. 10.3389/fcell.2022.1007614

Paim, L. M. G., & FitzHarris, G. (2022, Mar 22). Cell size and polarization determine cytokinesis furrow ingression dynamics in mouse embryos. Proc Natl Acad Sci U S A, 119(12), e2119381119. 10.1073/pnas.2119381119

Piekny, A. J., & Glotzer, M. (2008, Jan 8). Anillin is a scaffold protein that links RhoA, actin, and myosin during cytokinesis. Curr Biol, 18(1), 30–36. 10.1016/j.cub.2007.11.068

Piekny, A. J., & Maddox, A. S. (2010, Dec). The myriad roles of Anillin during cytokinesis. Semin Cell Dev Biol, 21(9), 881–891. 10.1016/j.semcdb.2010.08.002

Pollard, T. D., & O’Shaughnessy, B. (2019, Jun 20). Molecular Mechanism of Cytokinesis. Annu Rev Biochem, 88, 661–689. 10.1146/annurev-biochem-062917-012530

Reymann, A. C., Staniscia, F., Erzberger, A., Salbreux, G., & Grill, S. W. (2016, Oct 10). Cortical flow aligns actin filaments to form a furrow. Elife, 5. 10.7554/eLife.17807

Roberts, B., Haupt, A., Tucker, A., Grancharova, T., Arakaki, J., Fuqua, M. A., Nelson, A., Hookway, C., Ludmann, S. A., Mueller, I. A., Yang, R., Horwitz, R., Rafelski, S. M., & Gunawardane, R. N. (2017, Oct 15). Systematic gene tagging using CRISPR/Cas9 in human stem cells to illuminate cell organization. Mol Biol Cell, 28(21), 2854–2874. 10.1091/mbc.E17-03-0209

Roberts, B., Hendershott, M. C., Arakaki, J., Gerbin, K. A., Malik, H., Nelson, A., Gehring, J., Hookway, C., Ludmann, S. A., Yang, R., Haupt, A., Grancharova, T., Valencia, V., Fuqua, M. A., Tucker, A., Rafelski, S. M., & Gunawardane, R. N. (2019, May 14). Fluorescent Gene Tagging of Transcriptionally Silent Genes in hiPSCs. Stem Cell Reports, 12(5), 1145–1158. 10.1016/j.stemcr.2019.03.001

Rodrigues, N. T., Lekomtsev, S., Jananji, S., Kriston-Vizi, J., Hickson, G. R., & Baum, B. (2015, Aug 27). Kinetochore-localized PP1-Sds22 couples chromosome segregation to polar relaxation. Nature, 524(7566), 489–492. 10.1038/nature14496

Schimmel, J., Munoz-Subirana, N., Kool, H., van Schendel, R., van der Vlies, S., Kamp, J. A., de Vrij, F., Kushner, S. A., Smith, G. C. M., Boulton, S. J., & Tijsterman, M. (2023, Jan 25). Modulating mutational outcomes and improving precise gene editing at CRISPR-Cas9-induced breaks by chemical inhibition of end-joining pathways. Cell Rep, 42(2), 112019. 10.1016/j.celrep.2023.112019

Shi, Y., Inoue, H., Wu, J. C., & Yamanaka, S. (2017, Feb). Induced pluripotent stem cell technology: a decade of progress. Nat Rev Drug Discov, 16(2), 115–130. 10.1038/nrd.2016.245

Singh, A. M. (2019). An Efficient Protocol for Single-Cell Cloning Human Pluripotent Stem Cells. Front Cell Dev Biol, 7, 11. 10.3389/fcell.2019.00011

Skryabin, B. V., Kummerfeld, D. M., Gubar, L., Seeger, B., Kaiser, H., Stegemann, A., Roth, J., Meuth, S. G., Pavenstadt, H., Sherwood, J., Pap, T., Wedlich-Soldner, R., Sunderkotter, C., Schwartz, Y. B., Brosius, J., & Rozhdestvensky, T. S. (2020, Feb). Pervasive head-to-tail insertions of DNA templates mask desired CRISPR-Cas9-mediated genome editing events. Sci Adv, 6(7), eaax2941. 10.1126/sciadv.aax2941

Sobral, A. F., Chan, F. Y., Norman, M. J., Osorio, D. S., Dias, A. B., Ferreira, V., Barbosa, D. J., Cheerambathur, D., Gassmann, R., Belmonte, J. M., & Carvalho, A. X. (2021, Dec 20). Plastin and spectrin cooperate to stabilize the actomyosin cortex during cytokinesis. Curr Biol, 31(24), 5415–5428 e5410. 10.1016/j.cub.2021.09.055

Spira, F., Cuylen-Haering, S., Mehta, S., Samwer, M., Reversat, A., Verma, A., Oldenbourg, R., Sixt, M., & Gerlich, D. W. (2017, Nov 6). Cytokinesis in vertebrate cells initiates by contraction of an equatorial actomyosin network composed of randomly oriented filaments. Elife, 6. 10.7554/eLife.30867

Strebinger, D., Deluz, C., Friman, E. T., Govindan, S., Alber, A. B., & Suter, D. M. (2019, Sep). Endogenous fluctuations of OCT4 and SOX2 bias pluripotent cell fate decisions. Mol Syst Biol, 15(9), e9002. 10.15252/msb.20199002

Su, K. C., Takaki, T., & Petronczki, M. (2011, Dec 13). Targeting of the RhoGEF Ect2 to the equatorial membrane controls cleavage furrow formation during cytokinesis. Dev Cell, 21(6), 1104–1115. 10.1016/j.devcel.2011.11.003

Takahashi, K., Tanabe, K., Ohnuki, M., Narita, M., Ichisaka, T., Tomoda, K., & Yamanaka, S. (2007, Nov 30). Induction of pluripotent stem cells from adult human fibroblasts by defined factors. Cell, 131(5), 861–872. 10.1016/j.cell.2007.11.019

Tamura, R., Jiang, F., Xie, J., & Kamiyama, D. (2021, Feb 26). Multiplexed labeling of cellular proteins with split fluorescent protein tags. Commun Biol, 4(1), 257. 10.1038/s42003-021-01780-4

Thorvaldsdottir, H., Robinson, J. T., & Mesirov, J. P. (2013, Mar). Integrative Genomics Viewer (IGV): high-performance genomics data visualization and exploration. Brief Bioinform, 14(2), 178–192. 10.1093/bib/bbs017

Tristan, C. A., Hong, H., Jethmalani, Y., Chen, Y., Weber, C., Chu, P. H., Ryu, S., Jovanovic, V. M., Hur, I., Voss, T. C., Simeonov, A., & Singec, I. (2023, Jan). Efficient and safe single-cell cloning of human pluripotent stem cells using the CEPT cocktail. Nat Protoc, 18(1), 58–80. 10.1038/s41596-022-00753-z

Turac, G., Hindley, C. J., Thomas, R., Davis, J. A., Deleidi, M., Gasser, T., Karaoz, E., & Pruszak, J. (2013). Combined flow cytometric analysis of surface and intracellular antigens reveals surface molecule markers of human neuropoiesis. PLoS One, 8(6), e68519. 10.1371/journal.pone.0068519

van Oostende Triplet, C., Jaramillo Garcia, M., Haji Bik, H., Beaudet, D., & Piekny, A. (2014, Sep 1). Anillin interacts with microtubules and is part of the astral pathway that defines cortical domains. J Cell Sci, 127(Pt 17), 3699–3710. 10.1242/jcs.147504

Verma, N., Zhu, Z., & Huangfu, D. (2017). CRISPR/Cas-Mediated Knockin in Human Pluripotent Stem Cells. Methods Mol Biol, 1513, 119–140. 10.1007/978-1-4939-6539-7_9

Viana, M. P., Chen, J., Knijnenburg, T. A., Vasan, R., Yan, C., Arakaki, J. E., Bailey, M., Berry, B., Borensztejn, A., Brown, E. M., Carlson, S., Cass, J. A., Chaudhuri, B., Cordes Metzler, K. R., Coston, M. E., Crabtree, Z. J., Davidson, S., DeLizo, C. M., Dhaka, S., Dinh, S. Q., Do, T. P., Domingus, J., Donovan-Maiye, R. M., Ferrante, A. J., Foster, T. J., Frick, C. L., Fujioka, G., Fuqua, M. A., Gehring, J. L., Gerbin, K. A., Grancharova, T., Gregor, B. W., Harrylock, L. J., Haupt, A., Hendershott, M. C., Hookway, C., Horwitz, A. R., Hughes, H. C., Isaac, E. J., Johnson, G. R., Kim, B., Leonard, A. N., Leung, W. W., Lucas, J. J., Ludmann, S. A., Lyons, B. M., Malik, H., McGregor, R., Medrash, G. E., Meharry, S. L., Mitcham, K., Mueller, I. A., Murphy-Stevens, T. L., Nath, A., Nelson, A. M., Oluoch, S. A., Paleologu, L., Popiel, T. A., Riel-Mehan, M. M., Roberts, B., Schaefbauer, L. M., Schwarzl, M., Sherman, J., Slaton, S., Sluzewski, M. F., Smith, J. E., Sul, Y., Swain-Bowden, M. J., Tang, W. J., Thirstrup, D. J., Toloudis, D. M., Tucker, A. P., Valencia, V., Wiegraebe, W., Wijeratna, T., Yang, R., Zaunbrecher, R. J., Labitigan, R. L. D., Sanborn, A. L., Johnson, G. T., Gunawardane, R. N., Gaudreault, N., Theriot, J. A., & Rafelski, S. M. (2023, Jan). Integrated intracellular organization and its variations in human iPS cells. Nature, 613(7943), 345–354. 10.1038/s41586-022-05563-7

Wang, T., Wei, J. J., Sabatini, D. M., & Lander, E. S. (2014, Jan 3). Genetic screens in human cells using the CRISPR-Cas9 system. Science, 343(6166), 80–84. 10.1126/science.1246981

Watanabe, K., Ueno, M., Kamiya, D., Nishiyama, A., Matsumura, M., Wataya, T., Takahashi, J. B., Nishikawa, S., Nishikawa, S., Muguruma, K., & Sasai, Y. (2007, Jun). A ROCK inhibitor permits survival of dissociated human embryonic stem cells. Nat Biotechnol, 25(6), 681–686. 10.1038/nbt1310

Weigert, M., Schmidt, U., Boothe, T., Muller, A., Dibrov, A., Jain, A., Wilhelm, B., Schmidt, D., Broaddus, C., Culley, S., Rocha-Martins, M., Segovia-Miranda, F., Norden, C., Henriques, R., Zerial, M., Solimena, M., Rink, J., Tomancak, P., Royer, L., Jug, F., & Myers, E. W. (2018, Dec). Content-aware image restoration: pushing the limits of fluorescence microscopy. Nat Methods, 15(12), 1090–1097. 10.1038/s41592-018-0216-7

Whitford, W., Hawkins, V., Moodley, K. S., Grant, M. J., Lehnert, K., Snell, R. G., & Jacobsen, J. C. (2022, May 20). Proof of concept for multiplex amplicon sequencing for mutation identification using the MinION nanopore sequencer. Sci Rep, 12(1), 8572. 10.1038/s41598-022-12613-7

Ye, J., Coulouris, G., Zaretskaya, I., Cutcutache, I., Rozen, S., & Madden, T. L. (2012, Jun 18). Primer-BLAST: a tool to design target-specific primers for polymerase chain reaction. BMC Bioinformatics, 13, 134. 10.1186/1471-2105-13-134

Yonemura, S., Hirao-Minakuchi, K., & Nishimura, Y. (2004, May 1). Rho localization in cells and tissues. Exp Cell Res, 295(2), 300–314. 10.1016/j.yexcr.2004.01.005

Yuce, O., Piekny, A., & Glotzer, M. (2005, Aug 15). An ECT2-centralspindlin complex regulates the localization and function of RhoA. J Cell Biol, 170(4), 571–582. 10.1083/jcb.200501097

Zhou, S., Feng, S., Brown, D., & Huang, B. (2020). Improved yellow-green split fluorescent proteins for protein labeling and signal amplification. PLoS One, 15(11), e0242592. 10.1371/journal.pone.0242592

